# Phase variable glycosylation in non-typeable *Haemophilus influenzae*

**DOI:** 10.1101/744037

**Authors:** Danila Elango, Benjamin L. Schulz

**Affiliations:** School of Chemistry and Molecular Biosciences, The University of Queensland, St Lucia 4072, Queensland, Australia; Australian Infectious Diseases Research Centre, The University of Queensland, Brisbane 4072, Australia; Centre for Biopharmaceutical Innovation, Australian Institute of Bioengineering and Nanotechnology, The University of Queensland, St Lucia 4072, Queensland, Australia

## Abstract

Non-typeable *Haemophilus influenzae* (NTHi) is a leading cause of respiratory tract infections worldwide and continues to be a global health burden. Adhesion and colonisation of host cells are crucial steps in bacterial pathogenesis, and in many strains of NTHi interaction with the host is mediated by the high molecular weight adhesins HMW1A and HMW2A. These adhesins are *N*-glycoproteins which are modified by cytoplasmic glycosyltransferases HMW1C and HMW2C. Phase variation in the number of short sequence repeats in the promoters of *hmw1A* and *hmw2A* directly affects their expression. Here, we report the presence of similar variable repeat elements in the promoters of *hmw1C* and *hmw2C* in diverse NTHi isolates. In an *ex vivo* assay, we systematically altered substrate and glycosyltransferase expression and showed that both of these factors affected the site-specific efficiency of glycosylation on HMW-A. Glycosylation occupancy was incomplete at many sites, variable between sites, and generally lower close to the C-terminus of HMW-A. We investigated the causes of this variability. As HMW-C glycosylates HMW-A in the cytoplasm, we tested how secretion affected glycosylation on HMW-A and showed that retaining HMW-A in the cytoplasm indeed increased glycosylation occupancy across the full length of the protein. Site-directed mutagenesis showed that HMW-C had no inherent preference for glycosylating asparagines in NxS or NxT sequons. This work provides key insights into factors contributing to the heterogenous modifications of NTHi HMW-A adhesins, expands knowledge of NTHi population diversity and pathogenic capability, and is relevant to vaccine design for NTHi and related pathogens.

## Introduction

*Haemophilus influenzae* is a Gram-negative, non-motile coccobacillus that belongs to the Pasteurellaceae family. The bacterium is a commensal microbe that commonly resides within the upper airways of humans (1), but is also an opportunistic pathogen that can cause mild, acute, chronic, recurrent, localized or invasive diseases (2–5). It is mostly associated with upper and lower respiratory tract infections, otitis media, sinusitis, conjunctivitis, and chronic obstructive pulmonary disease (COPD) (5, 6). *H. influenzae* can be characterised into two distinct types: those encapsulated by a polysaccharide capsule and those that are unencapsulated, commonly termed non-typeable *H. influenzae* (NTHi) (2). There are six known capsular serotypes (serotypes a-f) of which serotype b (Hib) was once one of the primary causes of pneumonia, sepsis, epiglottitis, and bacterial meningitis (2, 7). While the incidence of invasive diseases caused by Hib has been largely controlled through the conjugated Hib vaccine (2, 8) (which is directed against type-specific polysaccharide capsule), this vaccine does not protect against NTHi, which is now an emerging pathogen and one of the leading causes of invasive bacterial disease (9). As a global health burden with neonates, young children, and the elderly as key vulnerable populations, there is a clear need to develop vaccines against NTHi (2, 10).

Efforts to develop a successful vaccine against NTHi are ongoing, with several antigenic virulence factors that contribute to the early stages of bacterial colonization under consideration as potential vaccine candidates (11–13). Several proteins are under investigation as vaccine candidates (11, 14), including fimbriae (OMP P5-homologous fimbriae (15)), outer membrane proteins (OMPs) (P2, P6) (16, 17), transferrin binding proteins (TbpB) (18), protein D (19), PilA (major subunit of type IV pili) (20), Hia (21), Hap (22), and HMW1/2 (14, 23). However, vaccine development is impeded by the high variability and heterogeneity of NTHi, especially of their surface molecules (13, 24). For example, strains vary in their adhesins (Hia, HMW, Hap) (24, 25), and there is high diversity in protein sequence between strains (24, 26). Adding further complexity, some genes undergo or are influenced by phase variable on/off expression (24, 27–29). This phase variation alters the expression of surface structures in a rapid and reversible manner, generating phenotypic heterogeneity in the bacterial population and ultimately changing the bacterium’s ability to adapt and survive in the host (29, 30).

Attachment to host epithelial cells is a crucial step in bacterial pathogenesis. In Gram-negative bacteria such as NTHi, autotransporter proteins are important virulence factors in the initial colonization of the host, including the high molecular weight adhesins HMW1 and HMW2, Hia (*H. influenzae* adhesin), or Hap (*Haemophilus* adherence and penetration protein) (25, 31). These adhesins are differentially distributed in *H. influenzae* isolates (25). Approximately 75% of clinical isolates possess the HMW adhesin system, while most of the remaining strains possess a Hia homolog that can still permit efficient attachment to cultured epithelial cells (25, 32–34). Furthermore, in strains lacking HMW or Hia, the Hap adhesin may allow for adherence to epithelial cells (32).

The NTHi HMW adhesin system (HMW-ABC) belongs to a family of two-partner secretion systems which consist of a secreted protein (TpsA; HMW1A) and a cognate outer membrane pore-forming translocator protein (TpsB; HMW1B) (35). In NTHi, the genes encoding the TPS system are within a three-gene locus; *hmw1ABC* or *hmw2ABC* (36, 37). HMW-A is a glycosylated adhesin protein, HMW-B is an outer membrane protein required for the secretion of the adhesin, and HMW-C is a soluble cytoplasmic glycosyltransferase that glycosylates HMW-A (38). The HMW1A (125kDa) and HMW2A (120kDa) adhesins are homologous and share 71% identity and 80% similarity (34), whereas HMW1B/HMW2B and HMW1C/HMW2C share 99% and 97% identity, respectively (39). The epithelial cell-binding domains (HMW1A: amino acids 555-914, and HMW2A: amino acids 553-916) are the parts of the proteins with maximal sequence difference (40). Indeed, HMW1A and HMW2A have different affinities to host cell glycan receptors which may drive cell or tissue tropism (41, 42).

The HMW-ABC *N*-glycosylation system of NTHi is unusual (Fig. 1). In most well-characterized systems *N*-glycosylation occurs in the eukaryotic endoplasmic reticulum or bacterial periplasm and requires an oligosaccharyltransferase (OST) to transfer a preassembled oligosaccharide from a lipid donor to asparagine residues in substrate proteins (43–45). In contrast, NTHi employs HMW-C, a soluble cytoplasmic protein belonging to the GT41 family of glycosyltransferases, a family which is otherwise comprised of *O*-GlcNAc transferases (46). HMW-C-like enzymes directly transfer hexose monosaccharides from UDP-hexose to acceptor proteins within the bacterial cytoplasm (47–49). Interestingly, both OST and HMW-C enzymes preferentially glycosylate Asn residues in glycosylation sequons (N-X-S/T; X≠P) (50). The *Actinobacillus pleuropneumoniae* HMW-C glycosyltransferase has been reported to catalyze both *N*- and *O*-linked glycosyltransferase reactions (51). NTHi HMW-A is modified with mono-hexose or di-hexose glycan structures, suggesting that HMW-C is also capable of forming hexose-hexose *O*-glycosidic bonds (45, 49, 51). Some bacteria, such *A. pleuropneumoniae*, contain ‘orphan’ HMW-Cs, in which *hmwC* and the gene encoding the target adhesin or autotransporter protein are at unlinked locations in the genome (52, 53). In NTHi, *hmwC* is adjacent to the *hmwAB* operon, although expression of *hmwAB* and *hmwC* are controlled by separate promoters. Furthermore, NTHi contains two homologous *hmw-ABC* loci (53): *hmw1ABC* and *hmw2ABC*.

**Figure 1.**
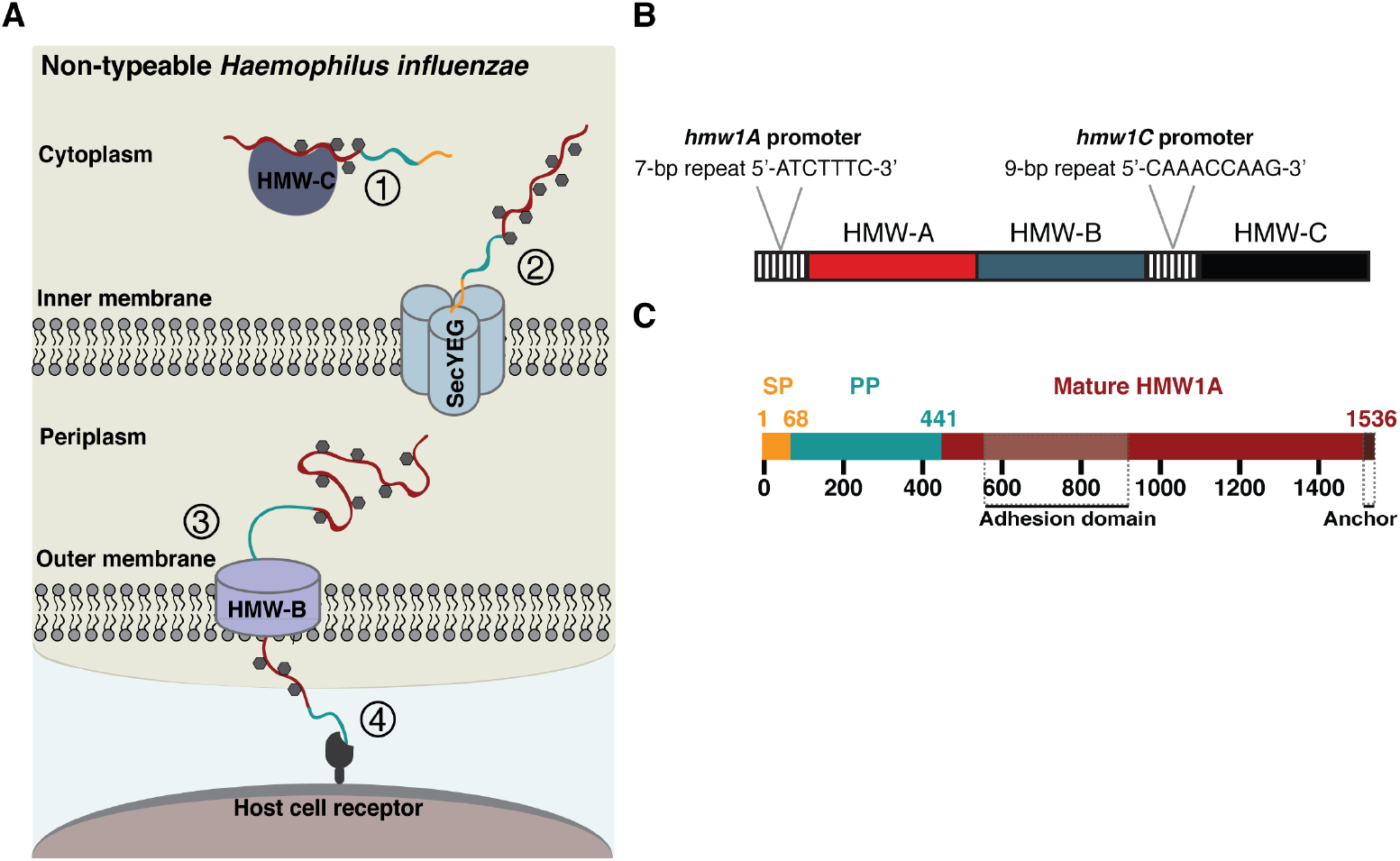
The *N*-glycosylation system of non-typeable *Haemophilus influenzae*. **(A)** The NTHi HMW-ABC two-partner secretion system. (1) HMW-A is translated, and glycosylated by HMW-C in the cytoplasm, then (2) translocated to the periplasm through the SecYEG translocon. (3) After translocation, the signal peptide is cleaved and the pro-protein interacts with the periplasmic domain of HMW-B (4). HMW-A is secreted across the outer membrane and retained on the bacterial cell surface via HMW-B. **(B)** The *hmw1A* promoter contains a 7-bp SSR (5’-ATCTTTC-3’) that regulates HMW1A expression (54). Another set of SSRs exist in the *hmw1C* promoter (5’-CAAACCAAG-3’). **(C)** HMW1A has three distinct regions: a signal peptide (amino acids 1-68), a pro-piece (HMW1A-PP) (amino acids 69-441), and mature HMW1A (amino acids 442-1536).

Genetic analysis of *hmw1A* and *hmw2A* has revealed that the expression of NTHi HMW-A is regulated by phase variation in the number of 7-bp short sequence repeats (SSRs) (5’-CTTTCAT-3’) within its promoter (54). This phase variation is thought to occur through slipped-strand mispairing and DNA polymerase slippage during replication of repetitive DNA sequences (54). The length of the promoter repeat tract is inversely correlated with the expression levels of HMW1A and HMW2A (54). As opposed to a typical “on/off” phase variable system, these promoter SSRs therefore allow for a gradient in HMW-A protein abundance and may contribute to heterogeneity in the bacterial population during infection (54). It is likely that phase variable expression of the HMW adhesins acts as an adaptive strategy in NTHi to aid in infection or evasion of host immune responses (55, 56). Specifically, population-level diversity may allow for selection of bacterial populations that best adapt to quickly-changing or niche environments and therefore allow for long-term survival within the human host.

Describing the full potential diversity of post-translational heterogeneity is critical for understanding the structure and function of the HMW-A adhesins and for their use as vaccine candidates. Here, we therefore assessed the effect of variable substrate (HMW-A) and glycosyltransferase (HMW-C) abundance on site-specific glycosylation of HMW-A, and investigated factors that control the efficiency of site-specific glycosylation in the adhesin.

## Methods

### Bacterial strain growth conditions

NTHi R2846 *hmw1AB* and *hmw1C* genes were used in this study. *Escherichia coli* cells expressing the gene/s of interest were grown at 37 °C in Luria-Bertani (LB) broth or agar plates (2% Bacto-tryptone, 1% Yeast extract, 2% NaCl). Antibiotics used to select for and maintain plasmids were ampicillin (100 μg/mL) and kanamycin (50 μg/mL). Various concentrations of arabinose and isopropyl β-D-1-thiogalactopyranoside (IPTG) were added to the media to induce protein expression where appropriate.

### Construction of plasmids

Oligonucleotide primers were designed to amplify DNA encoding *hmw1AB* incorporating *Nco*I and *Sac*I restriction sites. PCR amplicons were digested with *Nco*I and *Sac*I (New England Biolabs, Ipswich, Massachusetts), purified using the Sigma-Aldrich gel extraction kit, and ligated into *Nco*I and *Sac*I-digested pET28a(+). Cloning with *NcoI* caused a D_2_N substitution in HMW1A, which was reverted to the native sequence using site-directed mutagenesis (57). The pBad-HMW1C plasmid previously constructed by Gawthorne (2014) (58) was used for this study. Site-directed mutagenesis (57) was used to introduce an S_1046_T point mutation in HMW1A and to delete sequence encoding the HMW1A signal sequence (amino acids 1-68). Top10 *E. coli* cells transformants were selected by growth on LB plates with kanamycin.

### Experimental design and statistical rationale

Experiments were performed *ex vivo* using BL21 Rosetta cells containing the pBad-*hmw1C* plasmid (58). Cells were grown in LB media containing 100 μg/mL ampicillin, and incubated at 37 °C until the cells reached mid-log phase. The cells were then chemically transformed with the pET28a(+) plasmid harbouring the *hmw1AB* gene, with transformants selected by growth on LB agar containing ampicillin and kanamycin. A single colony was then inoculated into 20 mL LB with 100 μg/mL ampicillin and 50 μg/mL kanamycin, and grown until an OD_600_ of 0.30. Varying amounts of arabinose and IPTG were supplemented to the media, and cells were incubated with shaking at 37 °C. Cells were harvested at an OD_600_ of 1 by centrifugation and cell pellets frozen at −20 °C. In the titration assays, we varied expression of HMW1AB with a range of concentrations of IPTG (0.05mM, 0.1mM, 0.2mM, 0.5mM, or 1mM) with 0.2% arabinose, and of HMW1C with a range of concentrations of arabinose (0.00002%, 0.0002%, 0.002%, 0.02%, 0.2%, or 2%.) with 0.1mM IPTG. For analysis of variant HMW1A, samples were prepared in triplicate to allow quantification and statistical analyses.

### Whole cell extraction

Proteins were denatured and reduced by resuspending frozen cell pellets in 1 mL 6 M guanidinium chloride, 100 mM Tris HCl buffer pH 7.5, and 10 mM DTT and incubation at 30 °C for 30 min with shaking. Cysteines were alkylated by addition of acrylamide to a final concentration of 50 mM and incubation at 30 °C for 30 min with shaking. Proteins were precipitated by adding 20 μL of reduced/alkylated protein sample to 400 μL of 1:1 methanol: acetone and incubation at −20 °C for at least 4 h. Samples were centrifuged at 18,000 rcf for 10 min, the supernatant removed, centrifuged for 1 min, all supernatant removed, and the pellets air dried. Each pellet was resuspended in 100 μL 100 mM ammonium acetate with 1 μg trypsin and incubated at 37 °C for 16 h with shaking.

### Mass spectrometry

Peptides were desalted using C18 ZipTips (Millipore). LC-ESI-MS/MS analysis was performed using a Prominence nanoLC system (Shimadzu) and TripleTof 5600 instrument (SCIEX) as described previously (59). Approximately 2 or 0.4 μg peptides were injected for data dependent acquisition (DDA) or data independent acquisition (DIA; Sequential Window Acquisition of All THeoretical Mass Spectra, SWATH-MS) (60), respectively. Peptides were separated on a VYDAC EVEREST reversed-phase C18 HPLC column (300 Å pore size, 5 μm particle size, 150 μm i.d. x 150 mm) at a flow rate of 1 μl/min with a gradient of 10-60% buffer B over 45 min, with buffer A (1% acetonitrile and 0.1% formic acid) and buffer B (80% acetonitrile with 0.1% formic acid). LC parameters were identical for DDA and DIA, and DDA and DIA MS parameters were set as previously described (61, 62).

### Mass spectrometry data analysis

Peptides and proteins were identified using ProteinPilot 5.1 (SCIEX), searching against *E. coli* K12, NTHi R2846 HMW1A, HMW1B, and HMW1C, trypsin and common contaminants (4500 total proteins), with settings: Sample type, identification; Cysteine alkylation, acrylamide; Instrument, TripleTof 5600; Species, none; ID focus, biological modifications; Enzyme, trypsin; Search effort, thorough ID. False discovery rate analysis using ProteinPilot 5.1 (SCIEX) was performed on all searches. Peptides identified with greater than 99% confidence and with a local false discovery rate of less than 1% were included for further analysis. MS/MS fragmentation spectra and peak selection were manually inspected where required using PeakView 2.1 (SCIEX), with settings: Shared peptides, allowed; Peptide confidence threshold, 99%; False discovery rate, 1%; XIC extraction window, 6 min; XIC width 75 ppm. The mass spectrometry proteomics data have been deposited to the ProteomeXchange Consortium via the PRIDE (63) partner repository with the dataset identifier PXD015046. Normalised protein abundance was calculated as the ratio of the intensity of the protein of interest (HMW1A or HMW1C), to the sum of all proteins (HMW1A, HMW1B, HMW1C, trypsin, and all detected *E. coli* proteins) (64). Glycan occupancy was measured as the ratio of the intensity of the glycosylated peptide to the sum of the intensities of the glycosylated and unmodified peptides (59). Extracted ion chromatograms (XICs) were generated using PeakView (SCIEX).

### DNA sequence and protein structural analysis

Tandem Repeats Finder (65) was used to find and analyse short sequence repeats (SSRs) in the promoter regions of the genes encoding HMW1A, HMW1C, HMW2A, and HMW2C of NTHi R2846, 2019, NCTC8143, 86-028NP, and PittEE (ADO96128.1, AKA47280.1, CKH04147.1, AAX88733.1, ABQ98149.1; ADO96126.1, AKA47278.1, CKH04095.1, AAX88735.1, ABQ98147.1; ADO96470.1, AKA47663.1, CKH14369.1, AAX88269.1, ABQ98487.1; ADO96472.1, AKA47665.1, CKH14421.1, AAX88267.1, ABQ98489.1), accessed 5^th^ August 2019. Multiple sequence alignment was performed using MUSCLE (66). The structural context of *N*-glycosylation sites on HMW1A was predicted using Jalviewer (67) and the JPred protein secondary structure prediction server, and based on JNETPRED (68).

## Results

### Presence of a variable number of repeat elements in the promoters of hmwA and hmwC

Previous studies reported the presence of variable repeat elements upstream of *hmw1A* and *hmw2A* in NTHi (27, 29, 54). We investigated the presence and diversity in variable SSRs in the promoter regions of *hmw1/2ab* from five different strains: R2846, 2019, NCTC8143, 86-028NP, and PittEE. Multiple sequence alignment of the region immediately upstream of the *hmw1/2A* loci showed that the previously reported 7-bp SSR (5’-CATCTTT-3’(54), referred to here as 5’-ATCTTTC-3’) was present in all five strains (Fig. 2). However, the precise number of repeats varied between NTHi strains and between *hmw1A* and *hmw2A*, consistent with previous findings (27). This variation likely reflects rapid phase variation in repeat length rather than consistent differences between strains.

**Figure 2.**
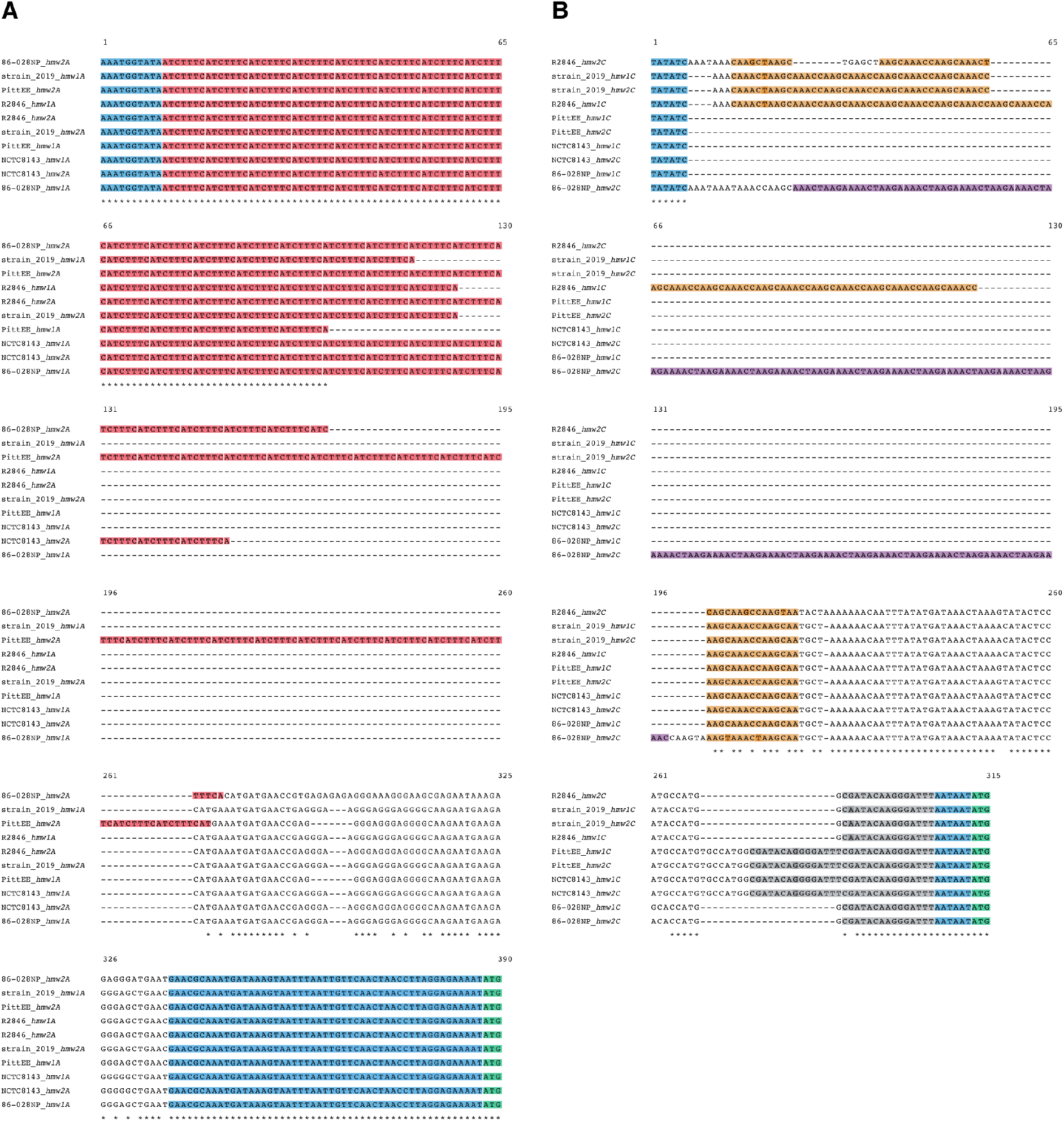
Variable number of repeat elements are present in the promoter regions of *hmw1A, hmw2A, hmw1C*, and *hmw2C* in diverse NTHi strains. Multiple sequence alignments of **(A)** *hmw1A* and *hmw2A* and **(B)** *hmw1C* and *hmw2C* promoter regions from strains R2846, 2019, NCTC8143, 86-028NP, and PittEE. Blue, conserved. Different colours highlight unique repeat sequences: *hmw-A* ‘ATCTTTC’, pink; *hmw-C* ‘CAAACCAAG’, yellow; *hmw-C* ‘AAACTAAGA’, purple; *hmw-C* ‘CGATACAAGGGATTT’, grey; start of open reading frame, green. Differences from the consensus repeat sequence are highlighted.

Inspection of the *hmw1C* and *hmw2C* loci in these same five strains also revealed the presence of repeat elements immediately upstream of *hmw1/2C*, but with different sequences to those observed at the *hmw1/2A* loci (Fig. 2). Out of the ten *hmw1C* and *hmw2C* promoter sequences inspected, eight contained repeat elements. The sequences upstream of R2846 *hmw1C* and strain 2019 *hmw1C* and *hmw2C* contained variable numbers of a 9-bp SSRs 5’-CAAACCAAG-3’. Strain 86-028NP had 19.4 copies of another repeat element 5’-AAACTAAGA-3’ preceding *hmw2C*. The sequence and the number of copies of the repeat elements also show some variation across different NTHi strains (Table 1). As variable repeat elements in *hmw-A* promoter regions affect HMW-A expression, these findings are consistent with similar variable length repeat elements in the *hmw1/2C* loci likely influencing *hmw1C* and *hmw2C* expression.

**Table 1.**
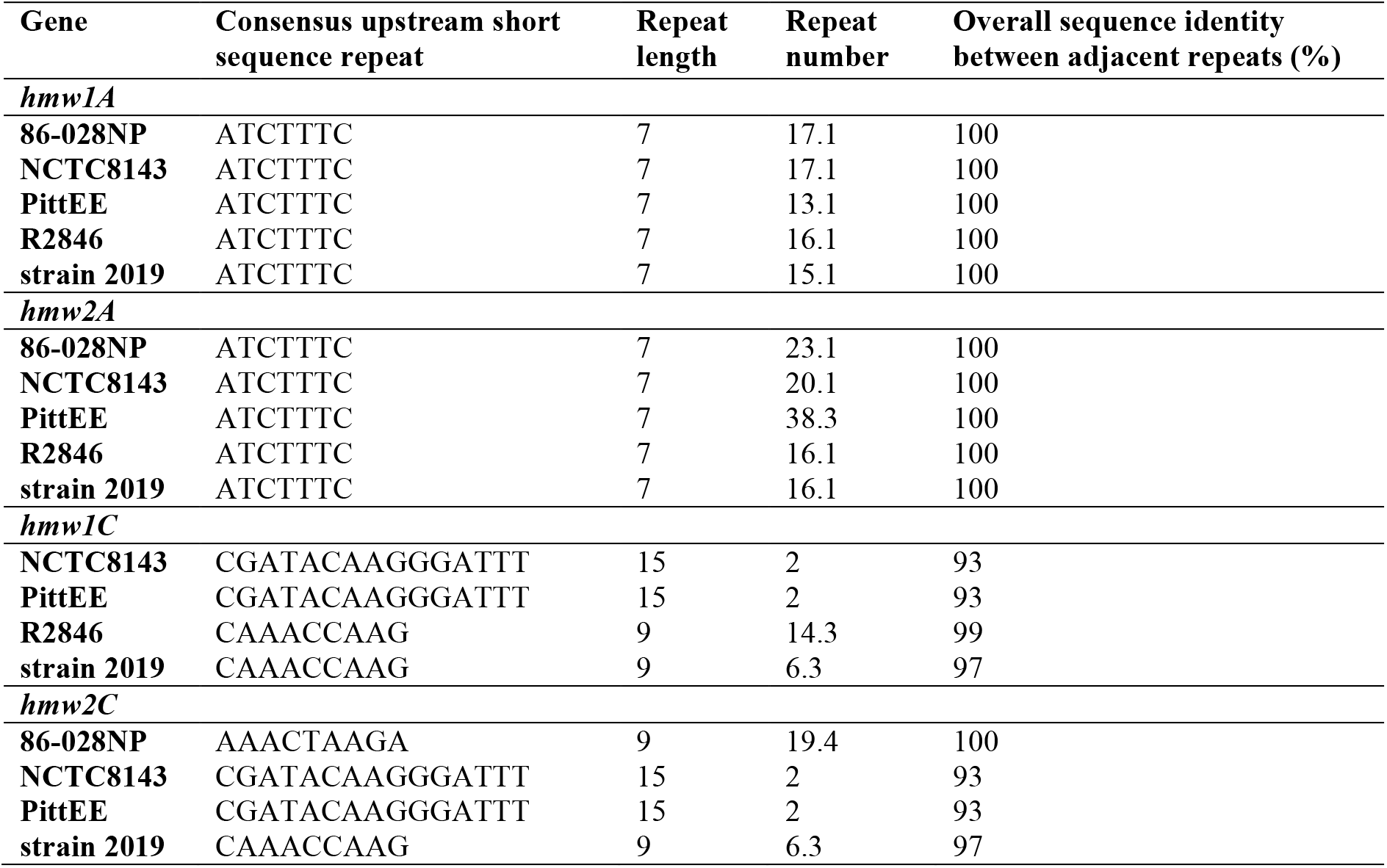
Short sequence repeats in the promoter regions of *hmw1A, hmw2A, hmw1C*, and *hmw2C* in different NTHi strains.

### A regulatable expression system for titration of HMW1A and HMW1C protein levels

Altered abundance of enzyme and substrate can affect the rate and quantitative extent of reactions. We therefore asked if changes in the relative abundance of the HMW-C glycosyltransferase and the HMW-A glycoprotein substrate affected site-specific glycosylation occupancy. Due to the inherent difficulty in controlling the expression of HMW1A and HMW1C *in vivo* in NTHi because of rapid phase variation in the length of repeat tracts, we tested the effect of variable substrate and glycosyltransferase abundance in an *ex vivo E. coli* system, similar to systems that have previously been used to study HMW-C activity (37, 58). Specifically, we cloned genes encoding the protein substrate (*hmw1AB*) and glycosyltransferase (*hmw1C*) into two separate plasmids and transformed both plasmids into BL21 *E. coli. hmw1AB* was cloned into the pET28 vector under the control of an IPTG-inducible promoter, and *hmw1C* was cloned into the pBAD vector under the control of an arabinose-inducible promoter. We then titrated the abundance of the proteins by supplementing the media with varying amounts of arabinose and IPTG, and performed DDA and DIA LC-ESI-MS/MS on whole cell extracts of *E. coli* expressing both HMW1A and HMW1C. With constant HMW1C abundance, controlled with 0.2% arabinose, we could increase the relative abundance of HMW1A by increasing the concentration of IPTG from 0.05 mM to 1 mM (Fig. 3A). The converse was also possible, keeping constant HMW1A abundance with 0.1 mM IPTG and increasing HMW1C abundance by increasing the concentration of arabinose from 0.00002% to 2% (Fig. 3B). This confirmed that the absolute and relative abundance of the HMW1C glycosyltransferase and its HMW1A glycoprotein substrate could be controlled in our heterologous expression system.

**Figure 3.**
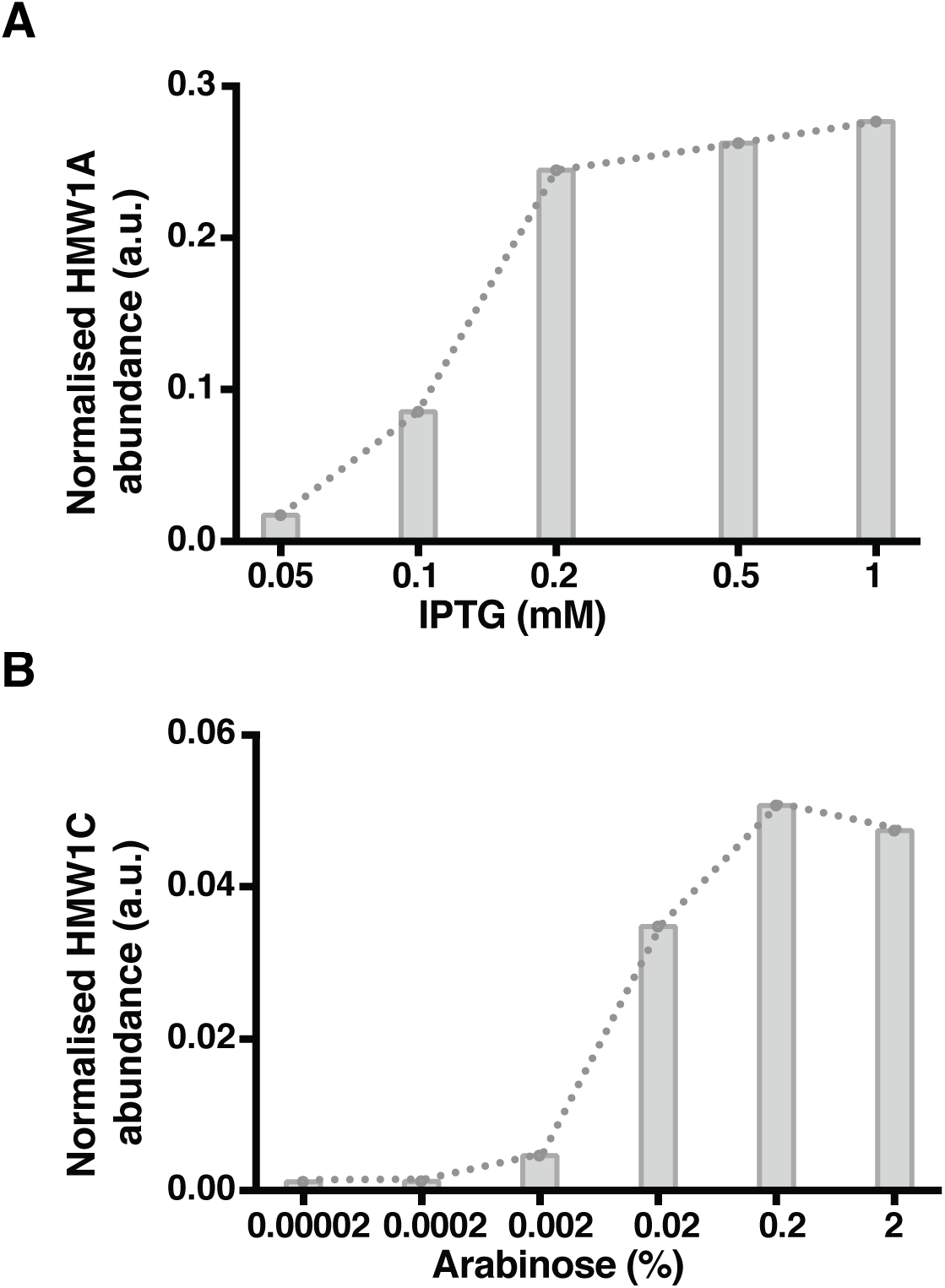
Titrated abundance of HMW1A and HMW1C. **(A)** Normalized abundance of HMW1A with 0.2% arabinose and varying concentrations of IPTG **(B)** Normalized abundance of HMW1C with 0.1mM IPTG and varying concentrations of arabinose.

### Site-specific glycosylation occupancy of HMW1A

We next performed a qualitative and quantitative assessment of glycosylated and unmodified tryptic peptides from HMW1A that were detected by DDA LC-ESI-MS/MS. We did not identify any peptides from the signal peptide region of HMW1A. Peptides belonging to the pro-piece (HMW1A-PP) were only identified in their non-glycosylated form, while numerous glycosylated and non-glycosylated peptides were identified from mature HMW1A. Coexpression of HMW1A and HMW1C allowed for robust detection of 45 sequon-containing peptides from HMW1A including 11 peptides with one sequon glycosylated with a single hexose and 19 unglycosylated peptides with one sequon (Table 2, Supplementary Figures S1-S18). In addition, we identified 8 distinct peptides and glycopeptides with more than one sequon. All glycosylation events were observed on Asn residues in sequons. Fig. 4 shows MS/MS spectra identifying tryptic peptides containing Asn484 in its glycosylated and unglycoyslated forms.

**Table 2.**
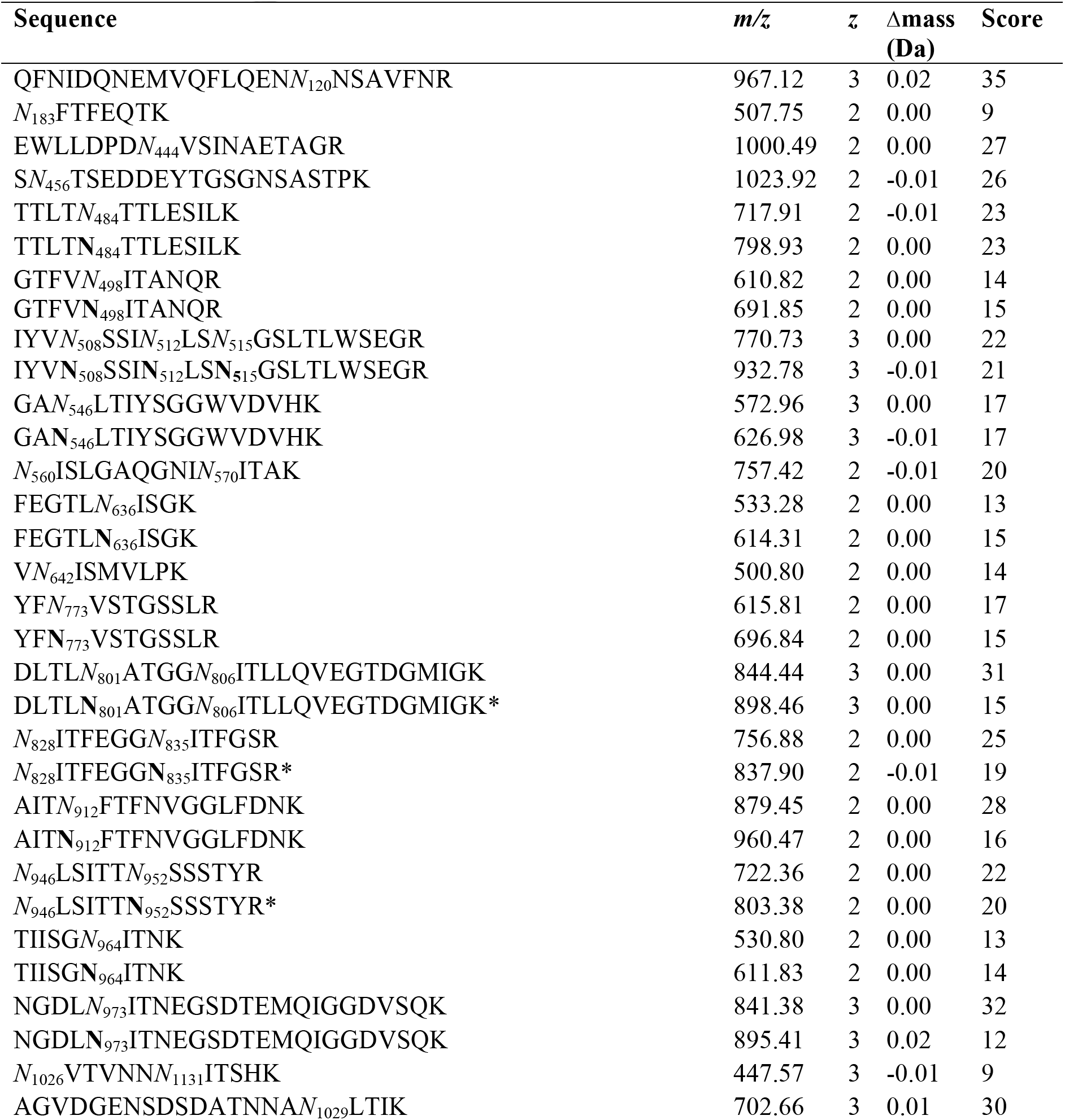
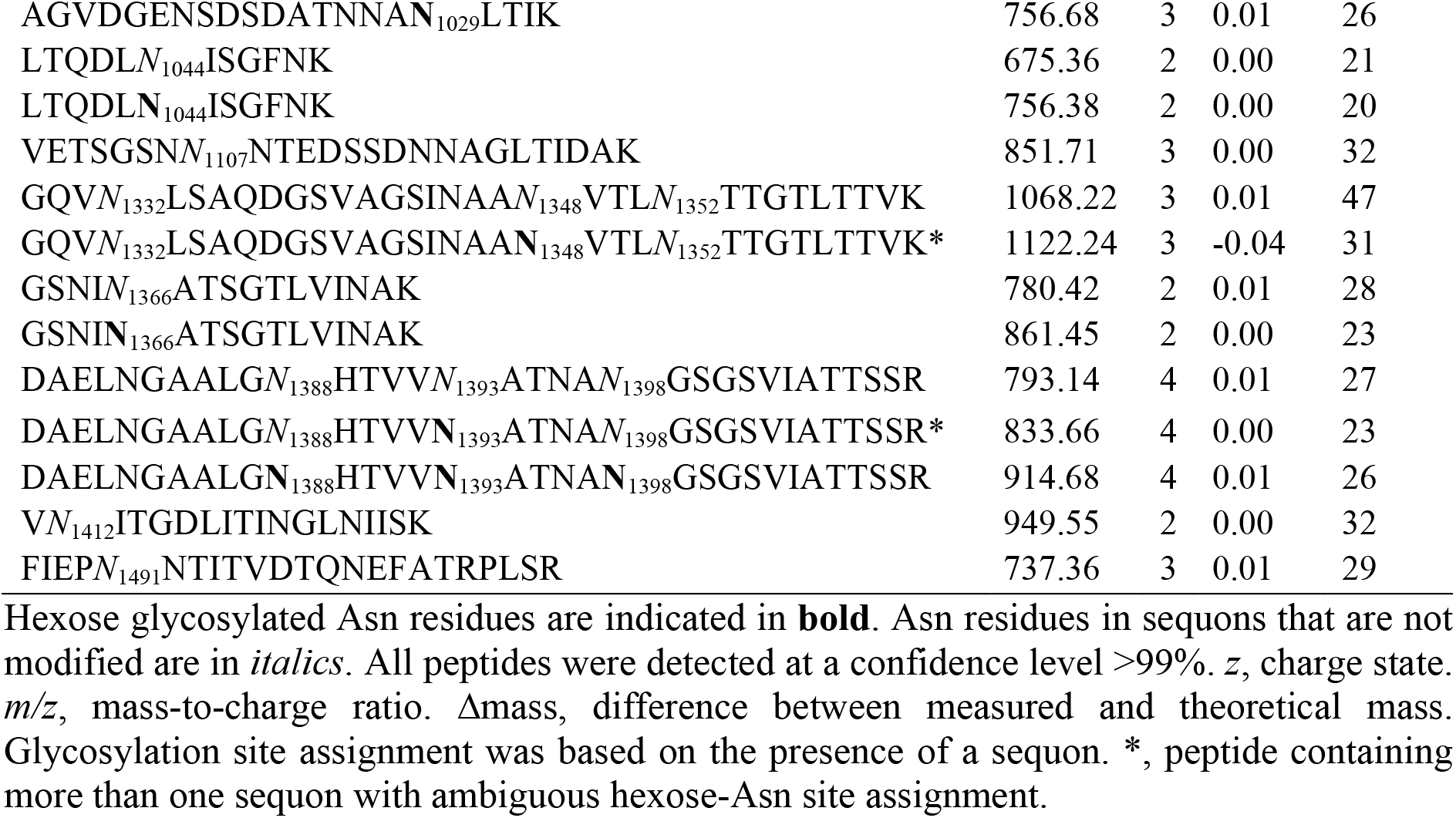
Glycosylated and non-glycosylated sequon-containing tryptic peptides from HMW1A identified by LC-ESI-MS/MS

**Figure 4.**
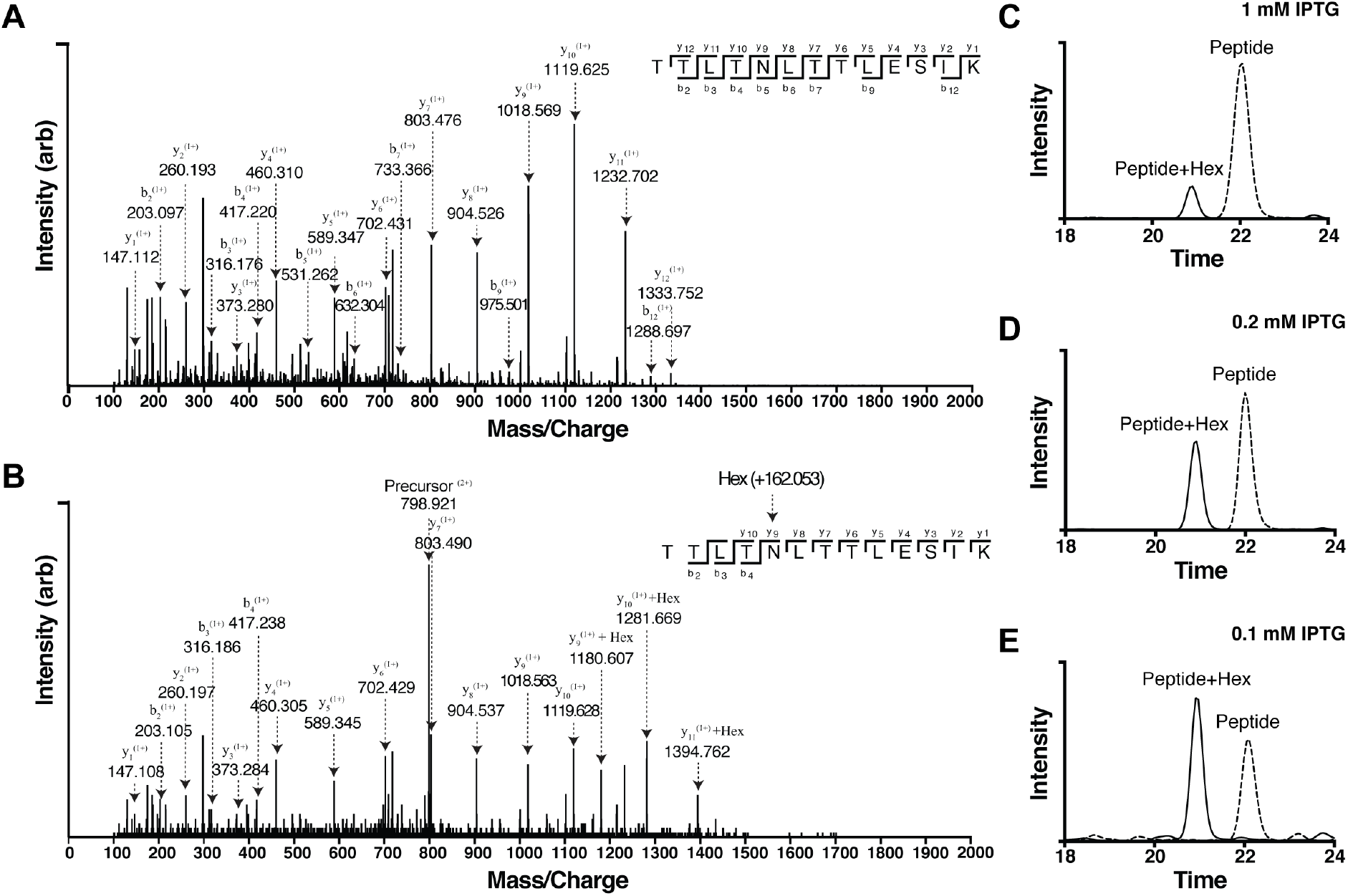
Identification and measurement of glycosylation occupancy on peptides containing HMW1A Asn484. **(A)** Annotated MS/MS spectra from DDA analysis of nonglycosylated peptide at an *m/z* of 717.9^2+^ and **(B)** glycopeptide at an *m/z* of 798.9^2+^ Not all matched fragment ions are labelled. Extracted ion chromatograms of glycosylated (solid) and non-glycosylated (dashed) peptides containing HMW1A Asn484 with HMW1C induced with 0.2% arabinose and HMW1A induced with **(C)** 1 mM IPTG, **(D)** 0.2 mM IPTG, or **(E)** 0.1 mM IPTG.

We next used SWATH/DIA detection of the unglycosylated and glycosylated forms of the same peptides to allow measurement of site-specific glycan occupancy across different *N*-glycosylation acceptor sequons in HMW1A. We measured site-specific glycan occupancy in peptides containing only a single *N*-glycosylation sequon. This is because of technical limitations in accurately assigning the site of modification in peptides with more than one *N*-glycosylation sequon. Inspecting one of the titration conditions (low expression of HMW1A (0.05mM IPTG) and high expression of HMW1C (0.2% arabinose), we found that there were quantitative differences in glycan occupancy between various sites in HMW1A. Most sites were efficiently glycosylated, but all were detected with at least some fraction unglycosylated; some had low occupancy (Asn912 and Asn1366); and others were completely unglycosylated (Asn120, Asn183, Asn444, Asn456, Asn642, Asn1107, Asn1412, and Asn1491). Glycosylation sites on bacterial *N*-glycoproteins are typically located in loops, as these structures are more flexible and are more accessible to glycosyltransferases (69, 70). We used the published structure of the HMW1A-PP (71) and the secondary structure of mature HMW1A predicted with JPred (68) to determine the location of glycosylated and nonglycosylated sequons in the secondary structure of HMW1A. HMW1A is mainly comprised of β sheets separated by short loops, and our data showed that glycosylated sequons were not any more likely to reside on a loop or secondary structural element than non-glycosylated sequons (Supplementary Figures S19 and S20).

### Titrating HMW1A substrate concentration affects site-specific glycan occupancy

Phase variation in the number of repeats in the promoter of *hmw1A* affects its expression (54). Using our *ex vivo* system, we therefore tested if changing HMW1A abundance also affected its site-specific glycosylation occupancy. We kept the abundance of the HMW-C glycosyltransferase fixed with 0.2% arabinose, controlled the abundance of HMW1A by addition of IPTG from 0.05 mM to 1 mM, and then used SWATH-MS to measure the site-specific glycosylation occupancy at 18 detectable sequons in HMW-A. This analysis showed that glycan occupancy in HMW-A was influenced by the concentration of the substrate protein in the cellular environment (Fig. 4 and 5).

**Figure 5.**
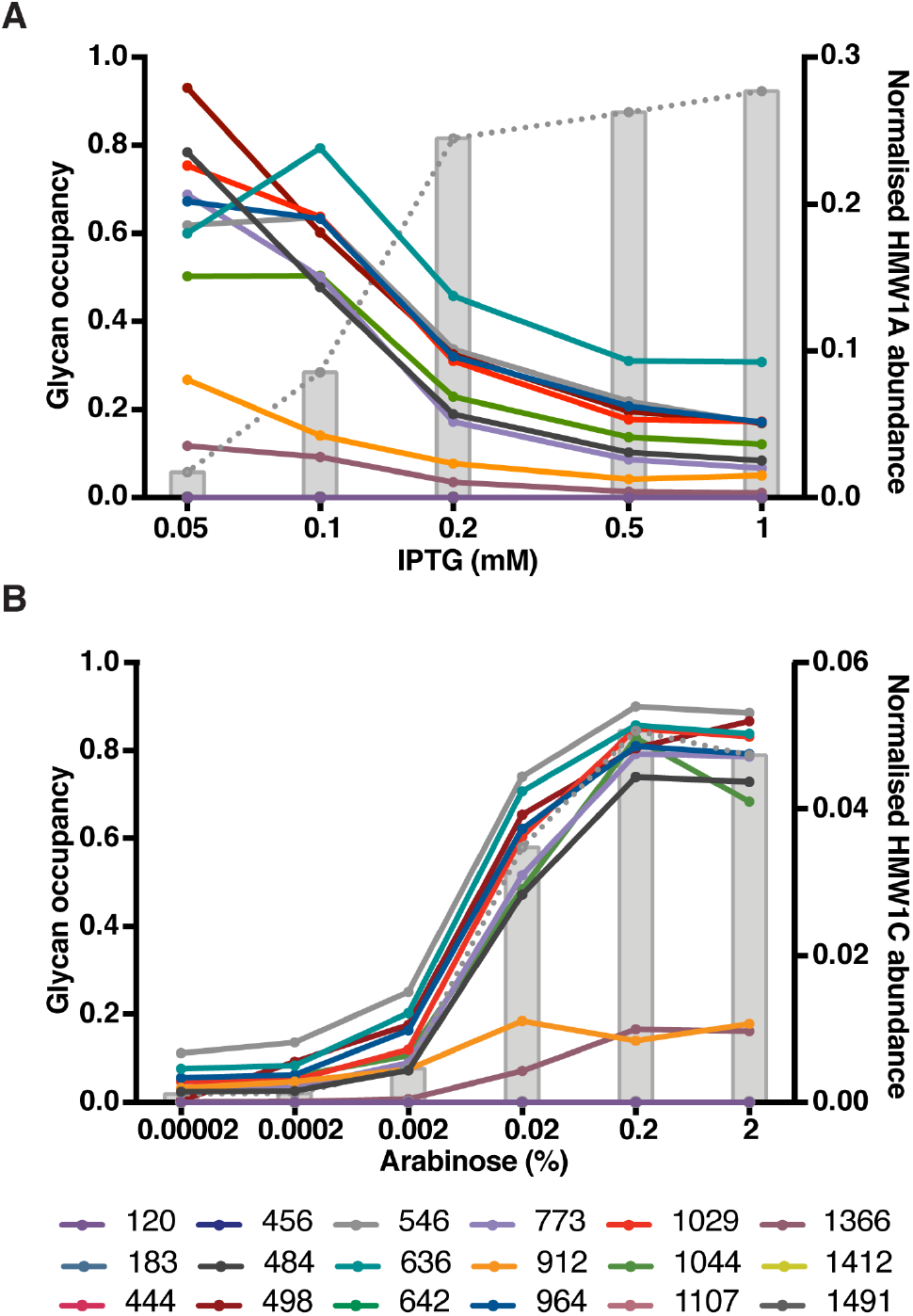
HMW1A and HMW1C abundance controls site-specific glycosylation occupancy on HMW1A. Site-specific glycosylation occupancy at 18 sites in HMW1A with **(A)** HMW1C abundance fixed with 0.2% arabinose and HMW1A abundance altered with varying concentrations of IPTG, and **(B)** HMW1A abundance fixed with 0.1 mM IPTG and HMW1C abundance altered with varying concentrations of arabinose.

We observed that as the abundance of the HMW1A protein substrate increased, site-specific glycan occupancy decreased at each site of modification in HMW1A (Fig. 5A). Our results were consistent with saturation of the HMW1C enzyme at high concentrations of HMW1A protein substrate. That is, at low HMW1A protein concentrations there was sufficient HMW1C enzyme to modify available sites in HMW1A, while at high expression levels of the HMW1A protein substrate the HMW1C enzyme had reduced capacity to modify acceptor sites. Our results therefore showed that titrating the expression of the HMW1A protein substrate could vary its site-specific extent of glycosylation.

### Titrating HMW1C glycosyltransferase concentration affects site-specific glycan occupancy on HMW1A

The presence and variation in the number of SSRs in the promoter region of *hmw1C* (Fig. 2B) suggested that HMW1C expression is influenced by the length of the repeat tract. We therefore studied the effect of variable HMW1C glycosyltransferase concentration on site-specific glycosylation occupancy of HMW1A. We varied HMW1C expression by addition of different amounts of arabinose (0.00002-2%), while the abundance of HMW1A was kept constant with 0.1 mM IPTG. We observed that varying the concentration of HMW1C resulted in changes in the extent of glycosylation across various sites in HMW1A (Fig. 5B). We observed a direct relationship between the concentration of the glycosyltransferase and glycosylation efficiency, whereby glycan occupancy in HMW1A increased with increasing HMW1C expression, until it asymptotically reached a point of saturation. This effect was consistent with an increased availability of glycosyltransferase increasing the efficiency of glycosylation of HMW1A. However, the maximum glycan occupancy in HMW1A (observed with 0.1 mM IPTG and 0.2% arabinose) did not reach greater than 90%, suggesting that factors apart from HMW1C abundance limited glycosylation efficiency.

### Glycosylation occupancy decreases towards the C-terminus of HMW1A

The pattern of glycosylation occupancy along the length of HMW1A was striking, with no glycosylation detected in HMW1A-PP, efficient glycosylation throughout most of the length of mature HMW1A, and lower occupancy glycosylation towards the C-terminus of the protein (Fig. 6). Decreased glycosylation efficiency close to the C-termini of proteins has also been reported for secretory eukaryotic glycoproteins (72, 73). While we observed a general decrease in site-specific glycan occupancy across HMW1A, the limited number of sequons we detected meant that the differences in glycosylation efficiency along the length of the protein did not reach statistical significance. We next investigated factors that might influence glycosylation efficiency at the C-terminus of HMW1A. The efficiency of eukaryotic co-translocational *N*-glycosylation decreases near the C-terminus of the substrate protein because polypeptide chain termination occurs before the acceptor site in the nascent polypeptide reaches the OST active site (73). Once translation of the polypeptide has been completed it is released from the ribosome and is therefore more rapidly translocated through the translocon and past the OST (73, 74). Although evolutionarily distinct from eukaryotic OST, a similar mechanism may be relevant to the HMW-ABC system. Glycosylation in NTHi occurs in the cytoplasm, and therefore once HMW-A is translocated into the periplasm it can no longer be glycosylated by HMW-C. It is therefore likely that secretion of HMW-A into the periplasm influences the glycosylation efficiency of HMW-A. Furthermore, the glycosylation of sequons located near the C-terminus of HMW-A would be more likely to be affected by this process. We therefore tested if HMW1A secretion affected the glycosylation efficiency of HMW1A. We created a variant of HMW1A lacking the residues corresponding to the signal peptide (amino acids 1-68), and co-expressed this truncated HMW1A together with HMW1C in BL21 *E. coli* cells with 0.1 mM IPTG and 0.2% arabinose. Whole cell extracts were prepared, tryptic digests analysed using LC-ESI-MS/MS, and SWATH-MS used to quantify glycosylation occupancy across HMW1A. Deletion of the HMW1A signal peptide did not specifically improve glycosylation at the C-terminus but rather globally increased glycosylation efficiency across all glycosylated Asn sites: nine out of ten partially modified sites were significantly more efficiently glycosylated (P<0.05, Student’s t-test) with deletion of the HMW1A signal peptide (Fig. 6A and B). All of the sequons that were not glycosylated in full length HMW1A remained unglycosylated in the variant lacking a signal peptide. Overall, our results suggested that when HMW1A is retained in the bacterial cytoplasm, glycosylation efficiency increases across the full length of the protein. This increase in efficiency may be due to increased exposure and interaction between the protein substrate and glycosyltransferase, and the absence of competition between glycosylation and secretion. These findings also suggest that although protein secretion limits glycan occupancy, this is not the cause of the inefficient glycosylation observed at the C-terminus of HMW1A, nor of the quantitative differences in glycosylation occupancy between different sequons throughout the protein.

**Figure 6.**
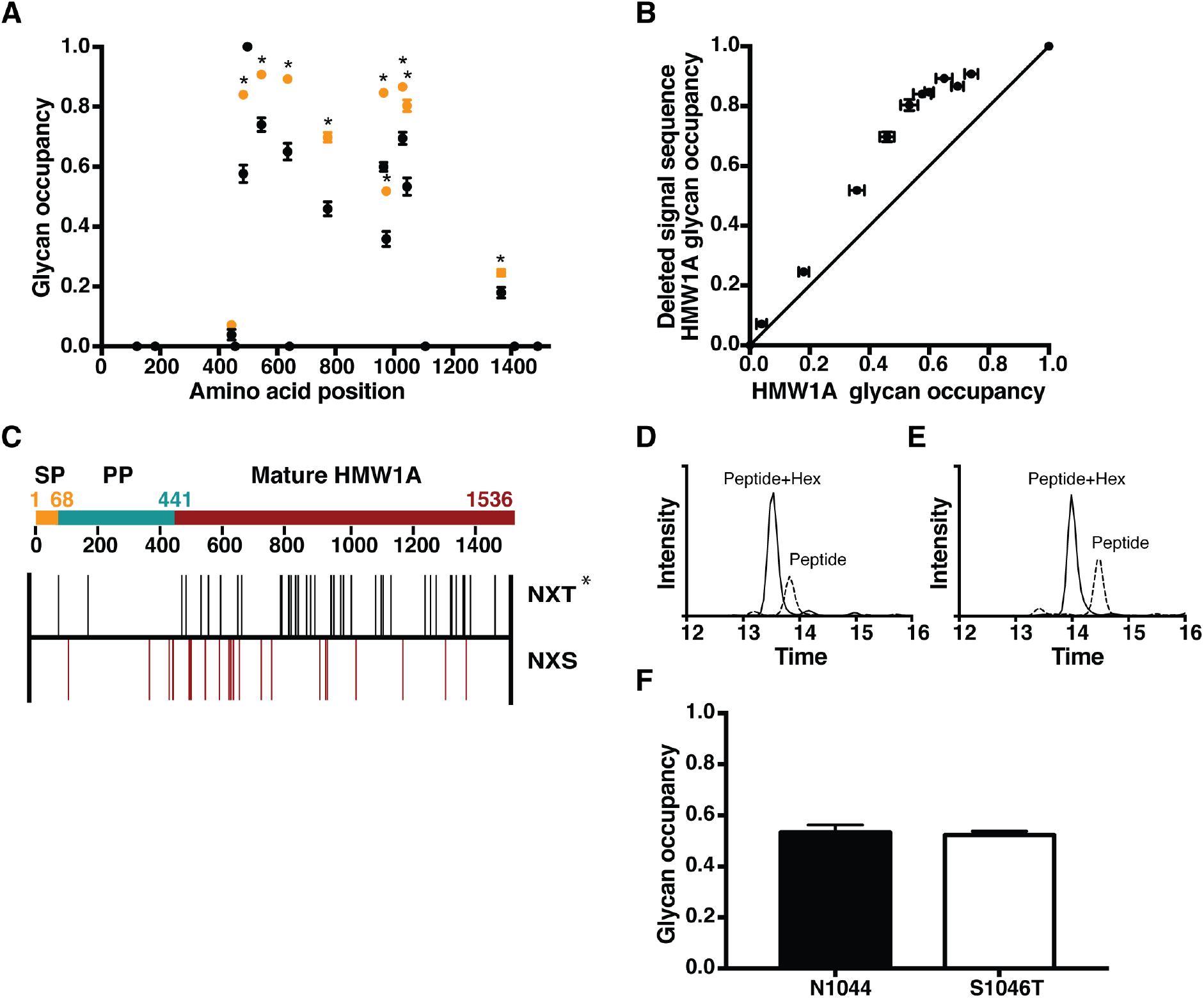
Factors affecting site-specific glycan occupancy across HMW1A. **(A)** Glycosylation site occupancy of Asn residues in HMW1A. • native HMW1A; 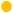 HMW1A with deleted signal peptide. Values are mean, error bars show standard error of the mean (n=3). *, significantly increased glycosylation site occupancy at these sites in HMW1A with deleted signal peptide compared with native HMW1A, P<0.05. **(B)** Site-specific glycan occupancy in deleted signal sequence HMW1A compared to full length HMW1A. Solid line indicates equal values. **(C)** Relative position of NXS (X≠P) and NXT (X≠P) sequons across HMW1A. *, difference in distribution of positions of NXS and NXT sequons in HMW1A was significantly different (P=0.011, two-tailed Mann-Whitney test). Extracted ion chromatograms of peptides and glycopeptides containing N_1044_ from **(D)** native (N_1044_IS_1046_) and **(E)** variant (S_1046_T) HMW1A. Dashed, peptide; solid, glycopeptide. **(E)** Glycan occupancy at N_1044_ in native and variant (S_1046_T) HMW1A. Values are mean. Error bars show standard error of the mean (n=3).

### HMW1C efficiently glycoslyates both NXS and NXT sequons

NXT sequons are more efficiently glycosylated than NXS sequons in a number of *N*-glycosylation systems (75), due to high affinity of Thr at the +2 position to the peptide binding site in OST (69). Upon inspection of HMW1A, we noted that there was an enrichment of NXT sequons and a depletion of NXS sequons towards the C-terminus of HMW1A (P-value= 0.011, Mann-Whitney) (Fig. 6C). This potentially correlated with the lower glycosylation occupancy we observed at sites close to the C-terminus of HMW1A (Fig. 6A). However, as occupancy was only measurable at relatively few *N*-glycosylation sites in HMW1A, it was difficult to statistically compare these features by observation alone. We therefore experimentally tested if NXS and NXT sequons in HMW1A were glycosylated with different efficiencies by HMW1C. We performed site-directed mutagenesis to convert Ser to Thr at amino acid 1046 (S_1046_T), changing the glycosylation sequon at N_1044_ from NIS to NIT. The plasmid (pET28-HMW1A_S1046T_B) was subsequently transformed into BL21/pBad-HMW1C cells. The variant HMW1A_S1046T_ and HMW1C were co-expressed with 0.1 mM IPTG and 0.2% arabinose, respectively. Control cells expressing HMW1C and wild type HMW1A with identical expression conditions were also prepared. The cells were harvested, and whole extracts prepared. Tryptic digests of the peptides were analysed using LC-ESI-MS/MS. We measured glycan occupancy at N_1044_ in native (Fig. 6D) and variant (Fig. 6E) HMW1A. This analysis showed that the mean glycosylation occupancy at N_1044_ in the mutant HMW1A was 53.3±2.9% (SEM) with NIS and 52.3±1.5% (SEM) with NIT, with no detectable significant difference (Fig. 6F). This suggested that the presence of serine or threonine at the +2 position of a sequon does not have a strong effect on its extent of glycosylation by HMW-C.

## Discussion

Expression of the HMW-A adhesin glycoprotein is highly variable in NTHi due to phase variation in the number of repeat elements in its promoter. Here, we established that this sequence variability is a feature of the promoter regions of *hmwA* and also of *hmwC* in diverse NTHi genomes (Fig. 2). This suggests that phase variable expression of *hmwA* and *hmwC* is a common feature of NTHi.

By independently titrating the expression of HMW1A and HMW1C in an *ex vivo* system, we showed that varying the abundance of the HMW1A glycoprotein substrate or of the HMW1C glycosyltransferase quantitatively affected site-specific glycosylation across HMW1A (Fig. 5). Site-specific glycosylation occupancy was increased either by increasing the abundance of the HMW1C glycosyltransferase, or by decreasing the abundance of the HMW1A glycoprotein. Phase variation in coding regions of genes that affects the presence or absence of the respective protein products is common in bacterial pathogens to enable immune evasion (76–78). Extending this paradigm, we propose that phase variation in the site-specific quantitative extent of glycosylation across the many glycosylation sites in HMW-A results in a similar diversification in proteoforms presented on the bacterial cell surface in a population of NTHi. This phase variability in the extent of glycosylation may then play a key role in immune evasion, and is also relevant for consideration of the use of HMW-A and other bacterial glycoproteins as vaccine candidates.

Even under conditions which provided the highest extent of glycosylation we could always detect some fraction of each site that was not modified. This suggested that factors other than the ratio of HMW1C glycosyltransferase to HMW1A glycoprotein substrate are important in controlling the extent of glycosylation in our *ex vivo* system. Amongst other potential factors, the extent of site-specific glycosylation depended on the position within HMW1A (Fig. 6). We identified peptides from HMW1A-PP and mature HMW1A, but no peptides from the signal peptide. This is consistent with rapid degradation of the signal peptide after cleavage by signal peptidase. When we investigated the structural context of sites that were efficiently and inefficiently modified, we found no difference in the localization of glycosylated and nonglycosylated sites on loop structures or secondary structural elements. The sequons located in the HMW1A-PP (at Asn120 and Asn183) were not glycosylated, but were also present in loops between β sheets. This suggests that aspects of local protein sequence or structure of the HMW1A-PP, or its accessibility to HMW1C, inhibited glycosylation in this region of the protein. It is also possible that lack of glycosylation of HMW1A-PP is critical for recognition by the HMW1B translocator pore or for translocation (79). This is plausible considering TpsA proteins contain a highly conserved ~250 amino acid TPS domain essential for protein secretion (79, 80).

We observed that glycosylation sites close to the C-terminus of HMW1A were generally inefficiently glycosylated (Fig. 6). When we deleted the signal sequence of HMW1A, the C-terminus of HMW1A remained relatively inefficiently glycosylated but there was a general increase in site-specific glycan occupancy across the full length of HMW1A (Fig. 6A and B). This indicated that secretion into the periplasm competes with glycosylation by HMW1C, but that this was not the underlying cause of inefficient glycosylation towards the C-terminus of HMW1A. We observed an enrichment of NXT sequons close to the C-terminus of HMW1A, perhaps evolved to increase the efficiency of glycosylation in this region of the protein (Fig. 6C). However, when we tested this by experimentally changing an NXS sequon to an NXT sequon in HMW1A, we found no significant difference in glycan occupancy between the native N_1044_IS_1046_ and variant N_1044_IT_1046_ (Fig. 6D-F). The relevance of inefficient glycosylation towards the HMW-A C-terminus remains unclear.

In summary, we have established that variable expression of HMW-A and HMW-C quantitatively influences glycosylation occupancy across diverse glycosylation sites in HMW-A. We predict that phase variable site-specific glycosylation facilitates antigenic escape or modulates adhesion by NTHi, and is therefore key in pathogenesis.

## Acknowledgements

We gratefully acknowledge the assistance of Dr Amanda Nouwens and Mr Peter Josh at The University of Queensland School of Chemistry and Molecular Biosciences Mass Spectrometry Facility, and the technical assistance of Toan Phung. BLS was funded by an Australian National Health and Medical Research Council RD Wright Biomedical (CDF Level 2) Fellowship APP1087975. This work was funded by an Australian Research Council Discovery Project DP160102766 and an Australian Research Council Industrial Transformation Training Centre IC160100027.

