## Supplementary Material for "Phase variable glycosylation in non-typeable *Haemophilus influenzae*"

**Supplementary Table 1. List of NTHi strains used in investigation of SSRs in the promoter regions of *hmw1A*, *hmw1C*, *hmw2A*, *hmw2C*. Genbank accessed 5th August 2019.**

| Strain | Genbank accession (Nucleotide) | Gene | GenBank accession (Protein) |
|---|---|---|---|
| **R2846/12** | CP002276.1 | *hmw1A* | ADO96128.1 |
| | | *hmw1C* | ADO96126.1 |
| | | *hmw2A* | ADO96470.1 |
| | | *hmw2C* | ADO96472.1 |
| **86-028NP** | CP000057.2 | *hmw1A* | AAX88733.1 |
| | | *hmw1C* | AAX88735.1 |
| | | *hmw2A* | AAX88269.1 |
| | | *hmw2C* | AAX88267.1 |
| **2019** | CP008740.1 | *hmw1A* | AKA47280.1 |
| | | *hmw1C* | AKA47278.1 |
| | | *hmw2A* | AKA47663.1 |
| | | *hmw2C* | AKA47665.1 |
| **NCTC8143** | LN831035.1 | *hmw1A* | CKH04147.1 |
| | | *hmw1C* | CKH04095.1 |
| | | *hmw2A* | CKH14369.1 |
| | | *hmw2C* | CKH14421.1 |
| **PittEE** | CP000671.1 | *hmw1A* | ABQ98149.1 |
| | | *hmw1C* | ABQ98147.1 |
| | | *hmw2A* | ABQ98487.1 |
| | | *hmw2C* | ABQ98489.1 |

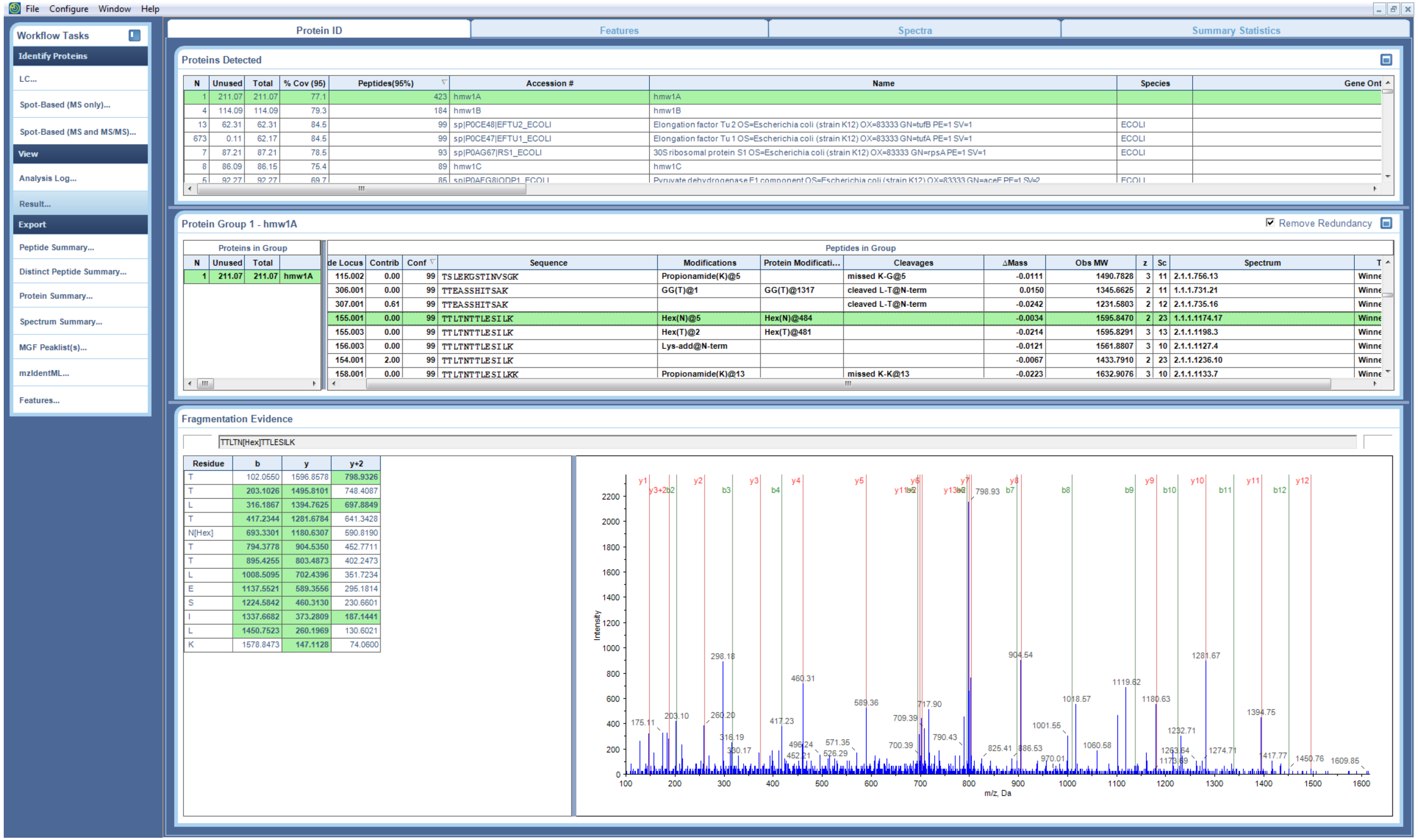


**Supplementary Figure S1. MS/MS peptide identification of TTLTN[+162.053]TTLESILK**

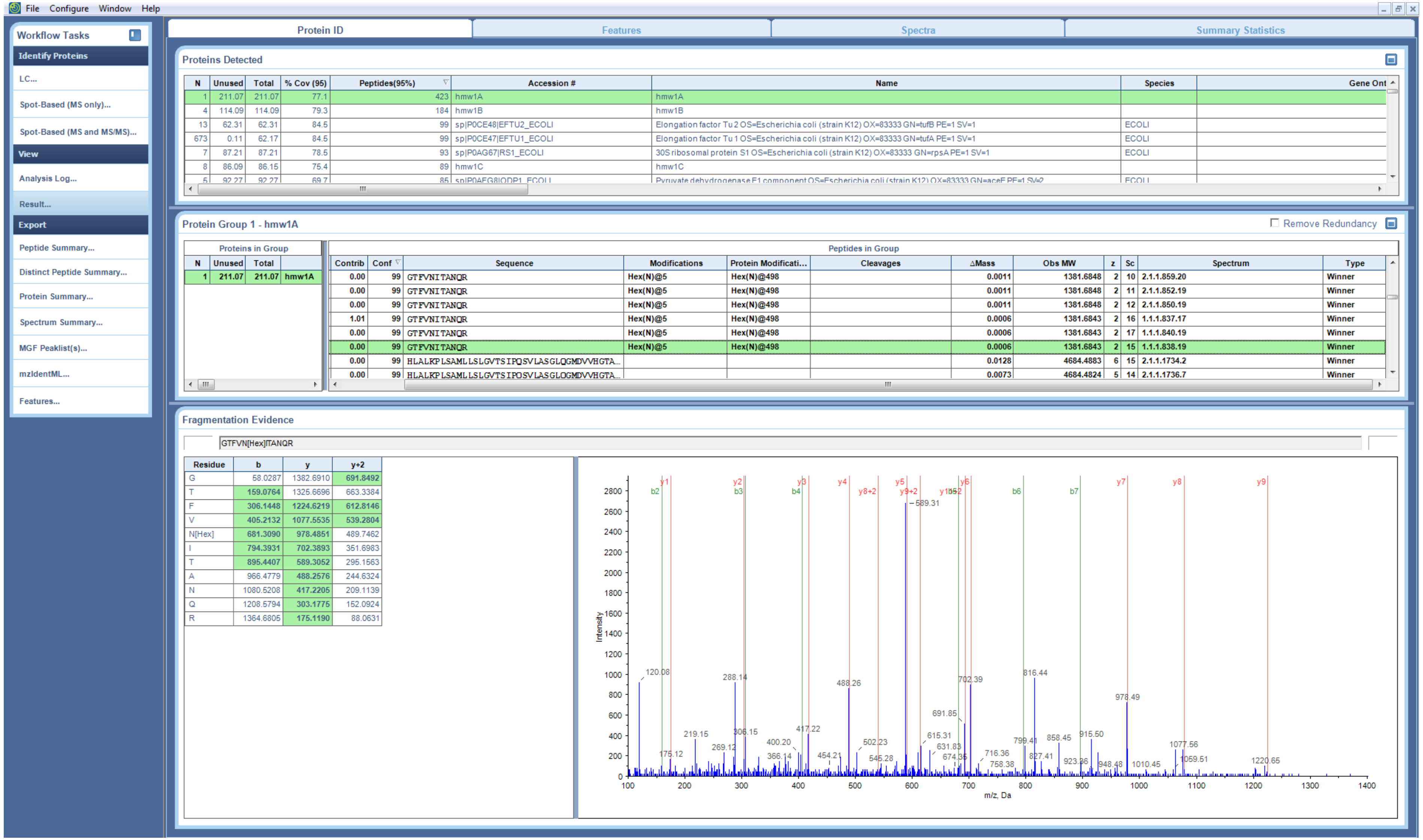


**Supplementary Figure S2. MS/MS peptide identification of GTFVN[+162.053]ITANQR**

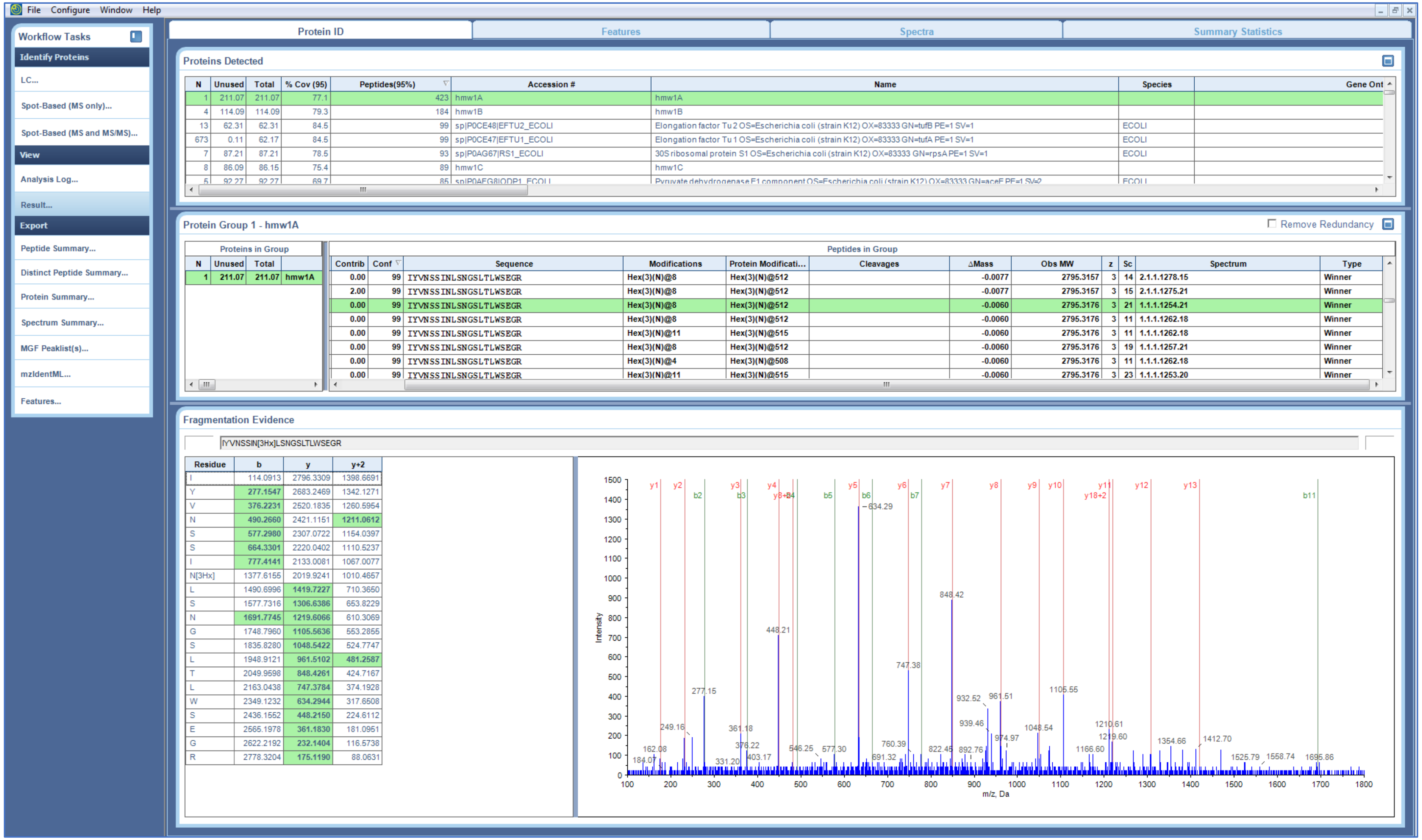


**Supplementary Figure S3. MS/MS peptide identification of IYVNSSIN[+486.159]LSNGSLTLWSEGR.** Site assignment of hexose-Asn is ambiguous, and assigned in Table 2 based on the presence of glycosylation sequons.

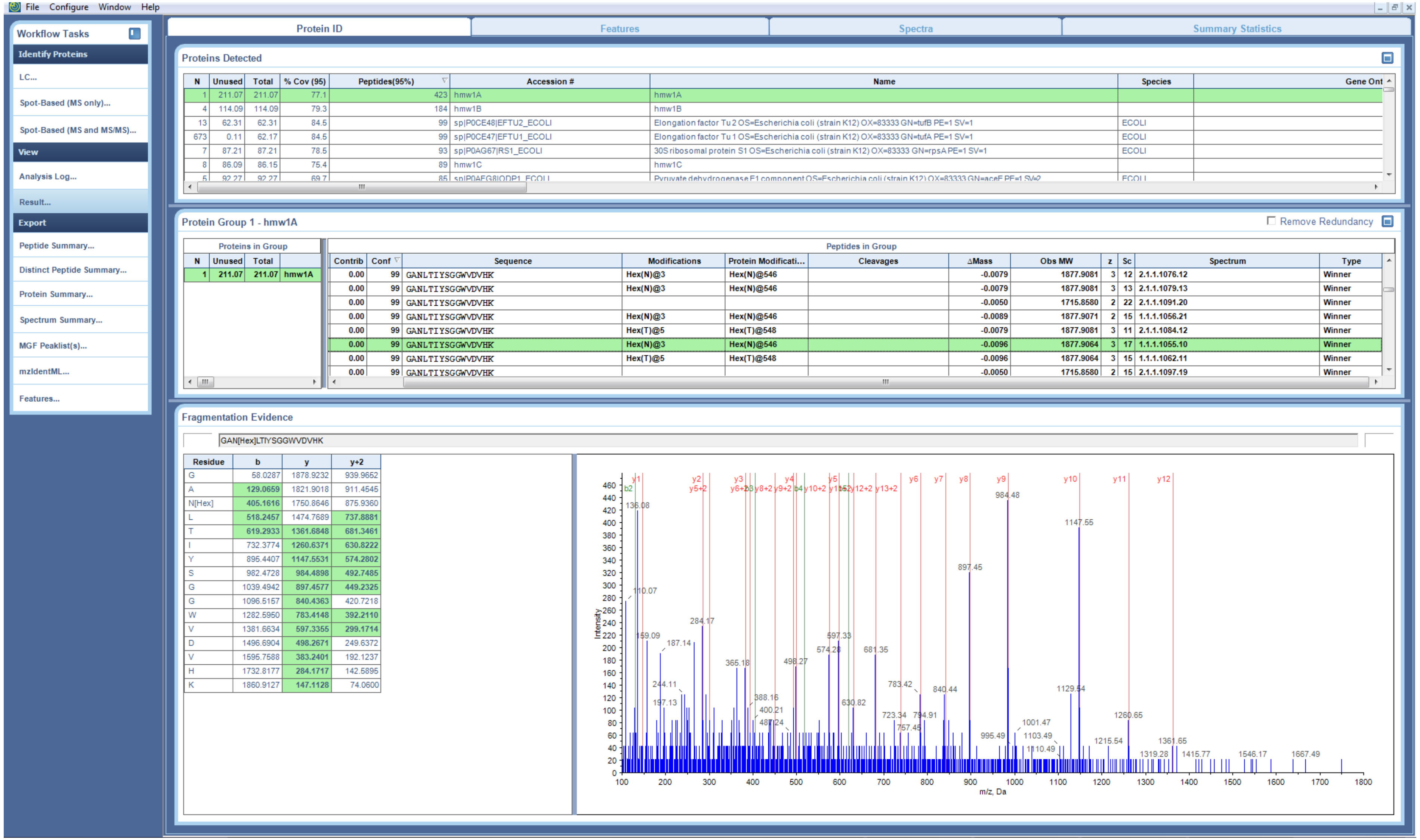


**Supplementary Figure S4. MS/MS peptide identification of GAN[+162.053]LTIYSGGWVDVHK**

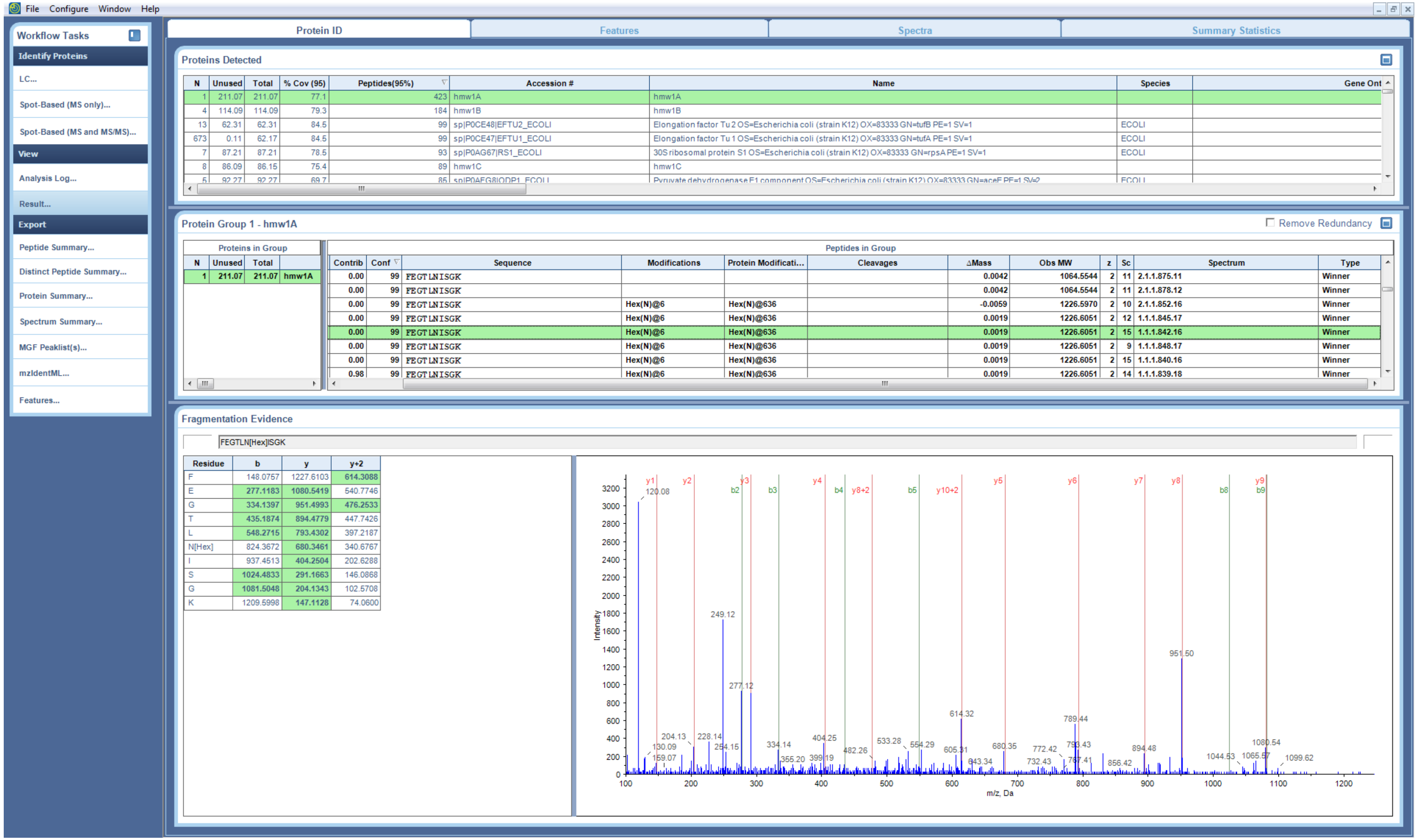


**Supplementary Figure S5. MS/MS peptide identification of FEGTLN[+162.053]ISGK**

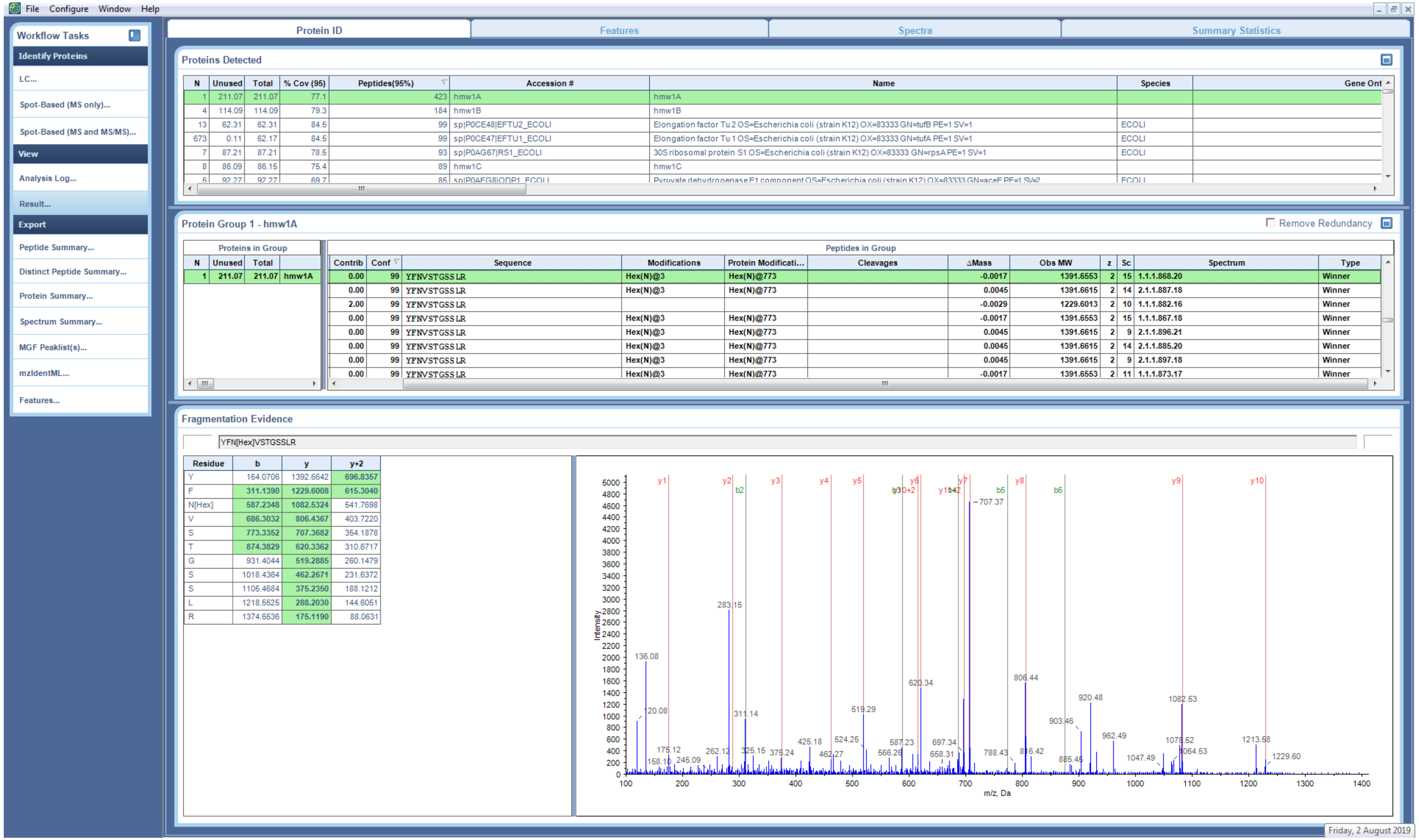


**Supplementary Figure S6. MS/MS peptide identification of YFN[+162.053]VSTGSSLR**

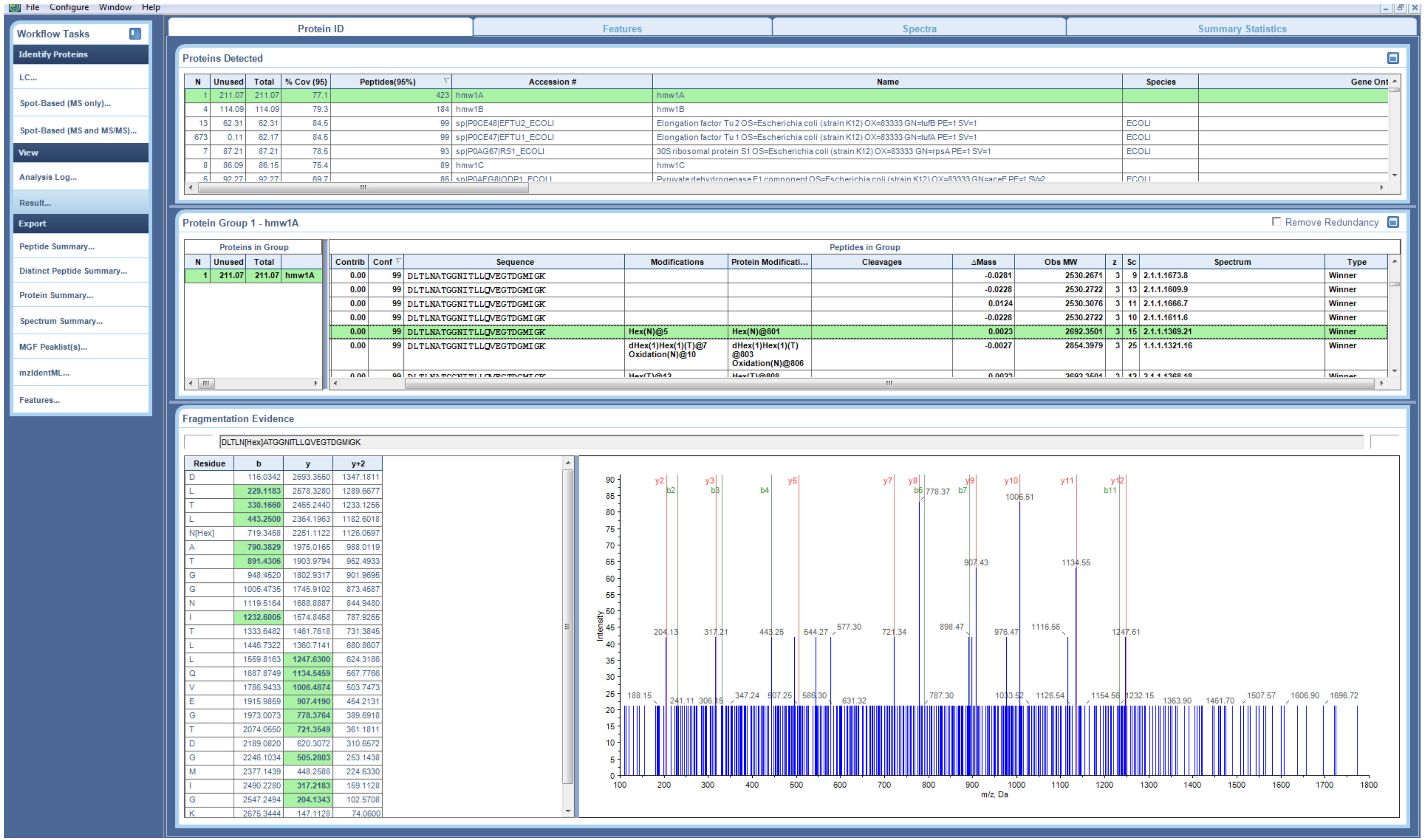


**Supplementary Figure S7. MS/MS peptide identification of DLTLN[+162.053]ATGGNITLLQVEGTDGMIGK**

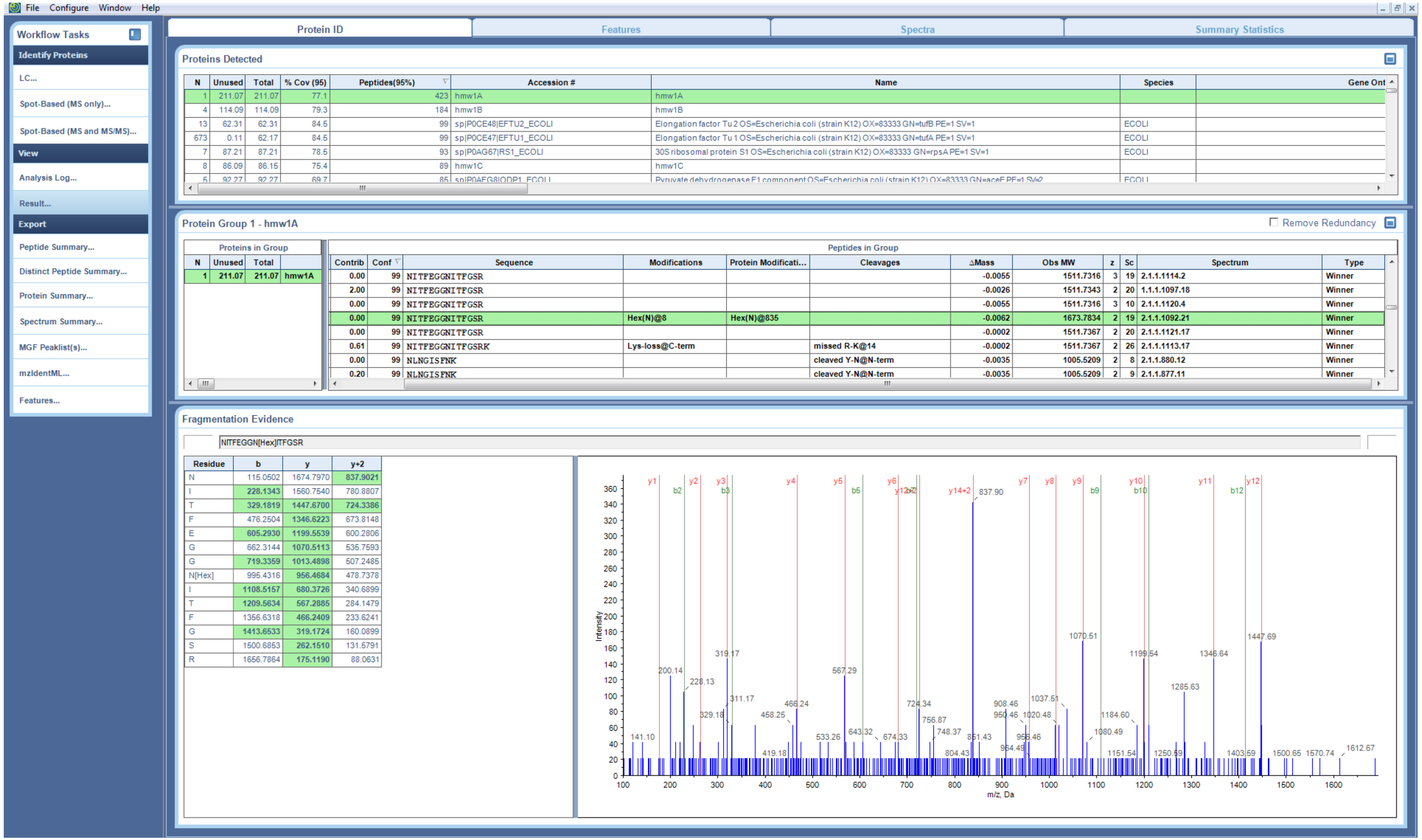


**Supplementary Figure S8. MS/MS peptide identification of NITFEGGN[+162.053]ITFGSR**

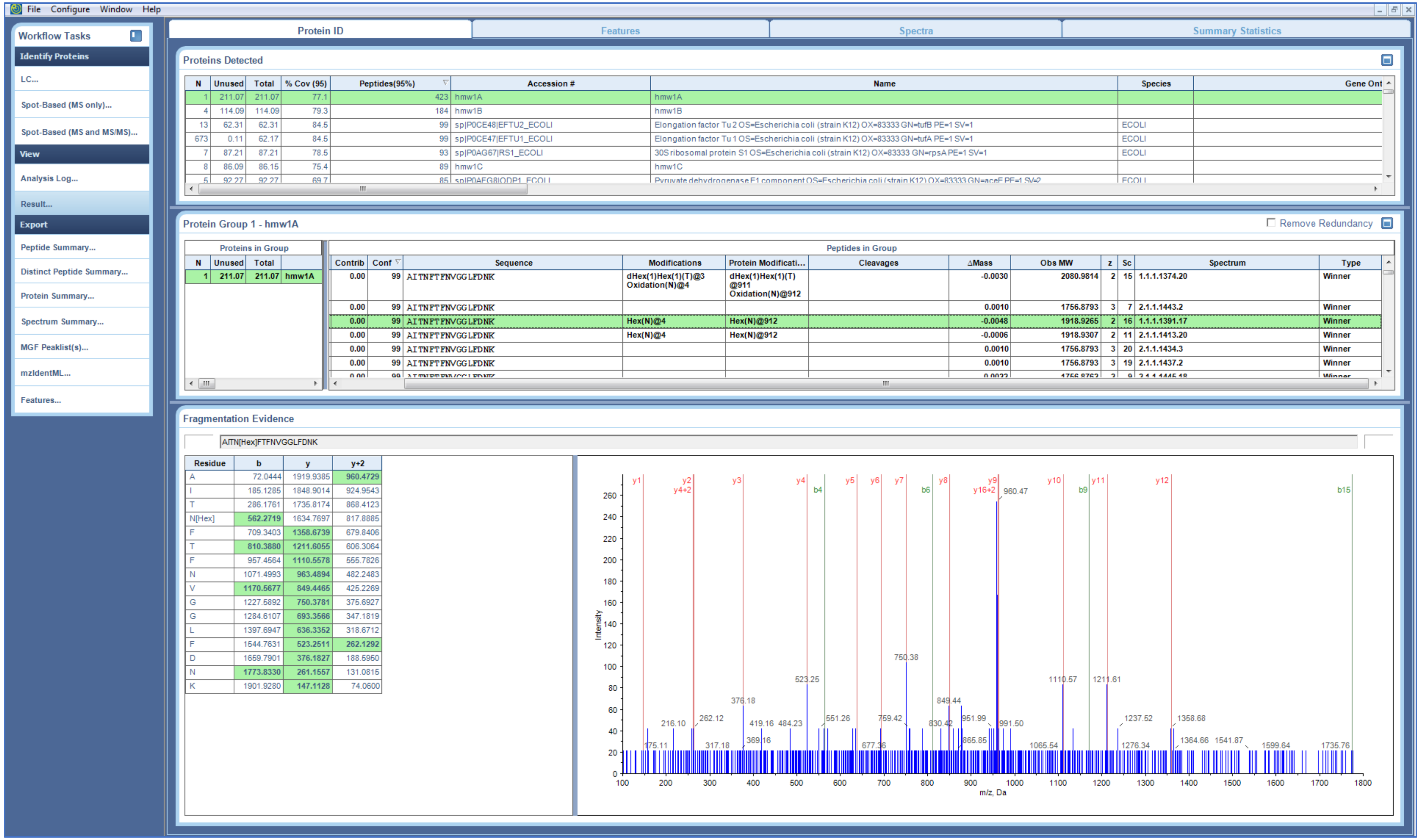


**Supplementary Figure S9. MS/MS peptide identification of AITN[+162.053]FTFNVGGLFDNK**

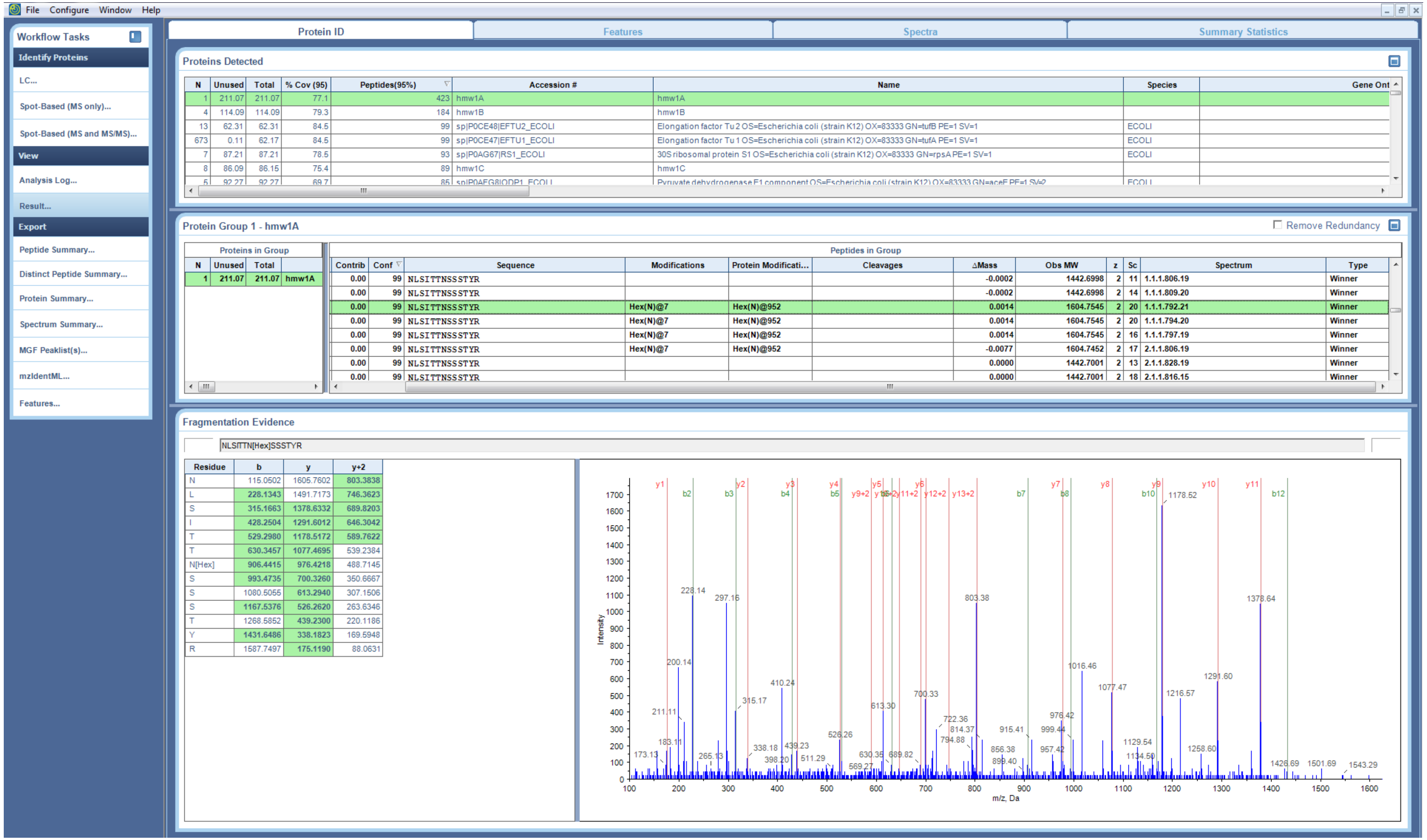


Supplementary Figure S10. MS/MS peptide identification of NLSITTN[+162.053]SSSTYR

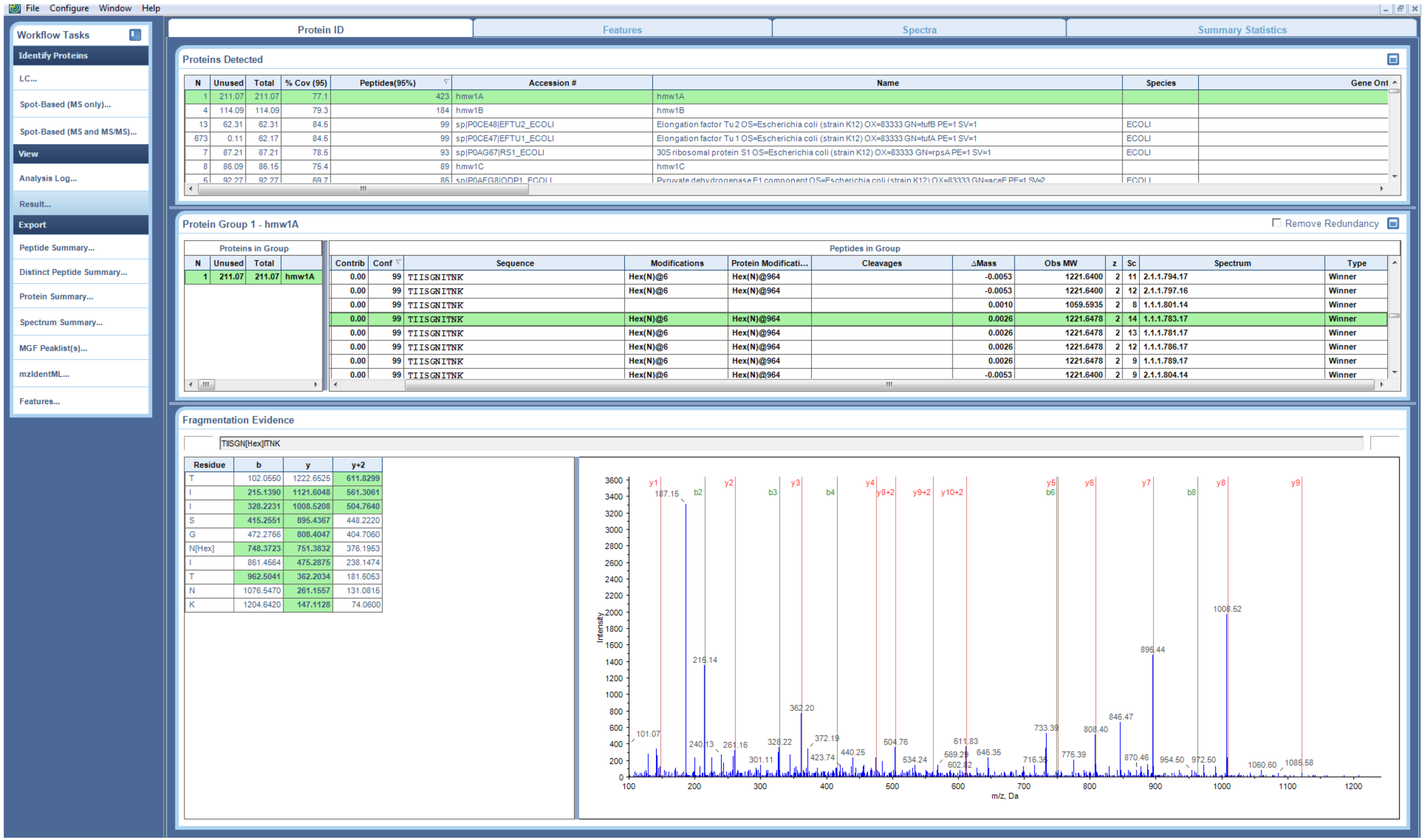


Supplementary Figure S11. MS/MS peptide identification of TIISGN[+162.053]ITNK

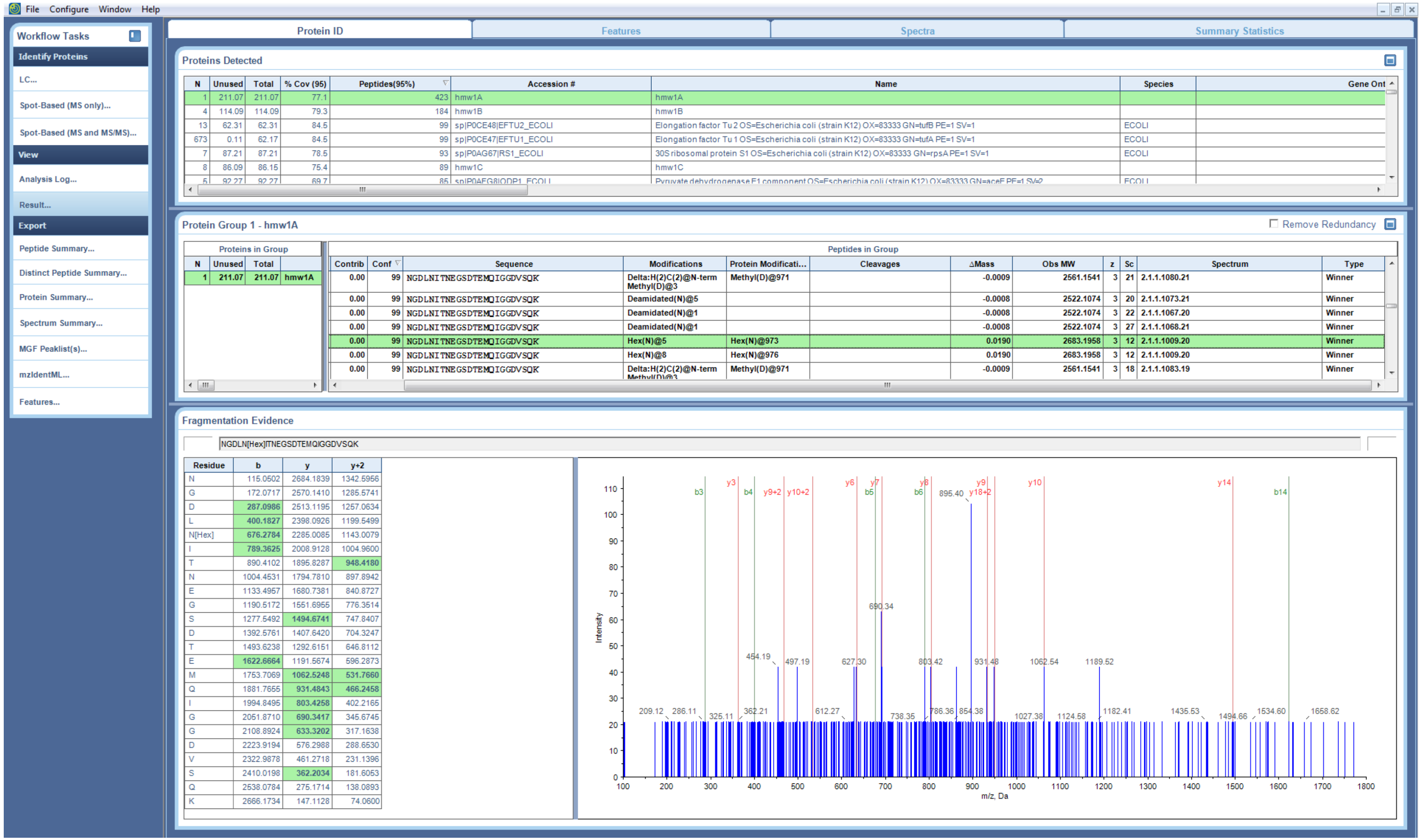


**Supplementary Figure S12. MS/MS peptide identification of NGDLN[+162.053]ITNEGSDTEMQIGGDVSQK**

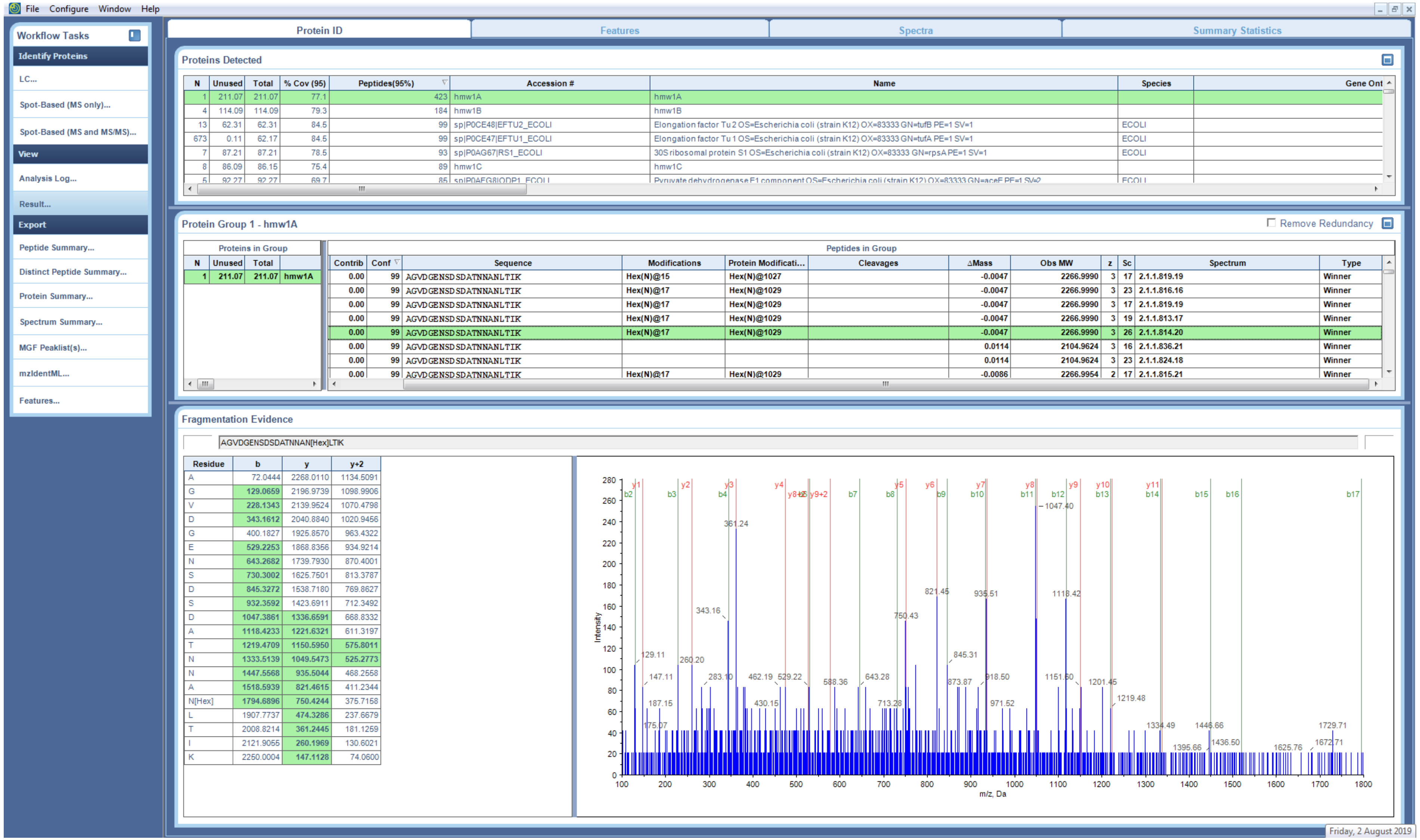


**Supplementary Figure S13. MS/MS peptide identification of AGVDGENSDSDATNNAN[+162.053]LTIK**

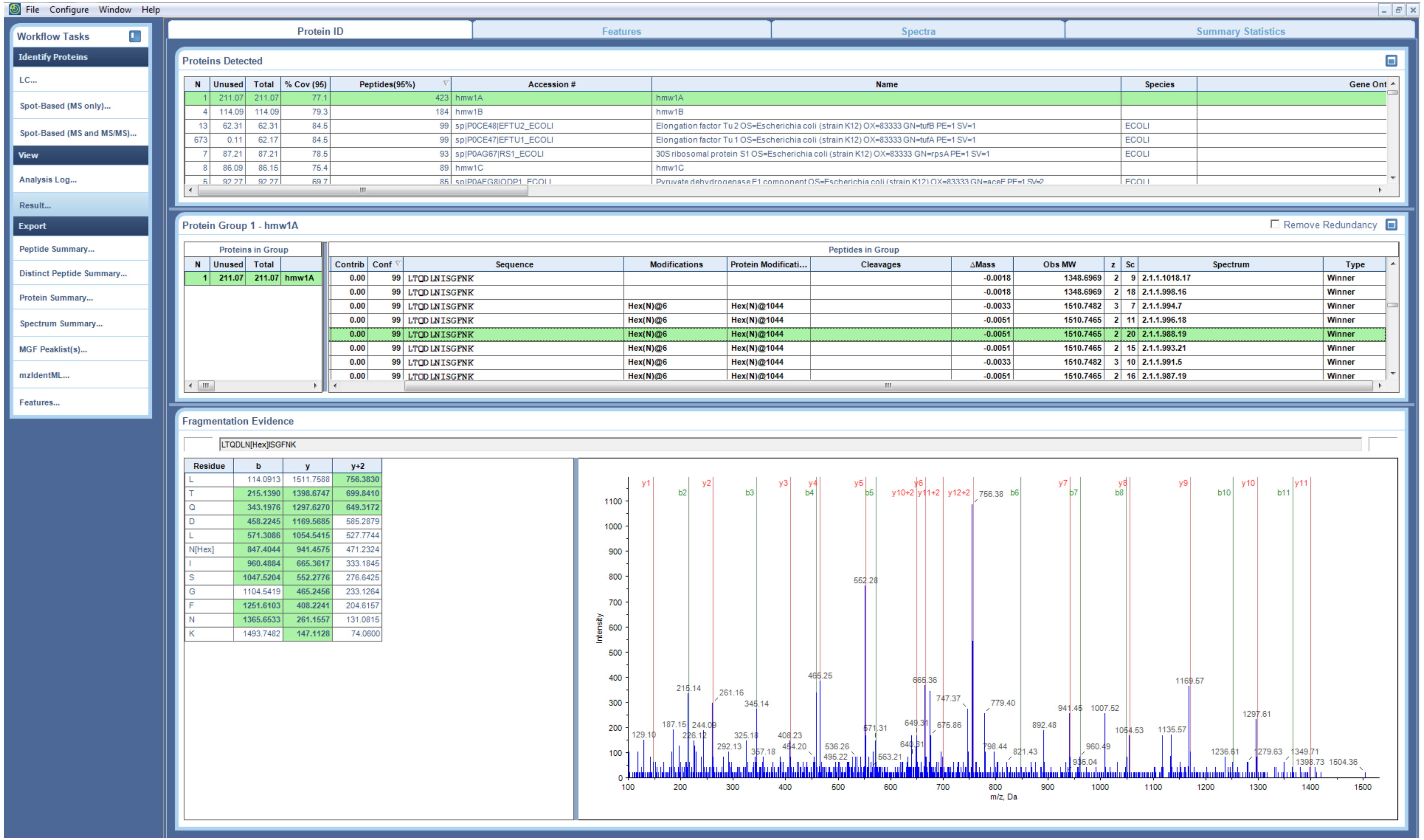


**Supplementary Figure S14. MS/MS peptide identification of LTQDLN[+162.053]ISGFNK**

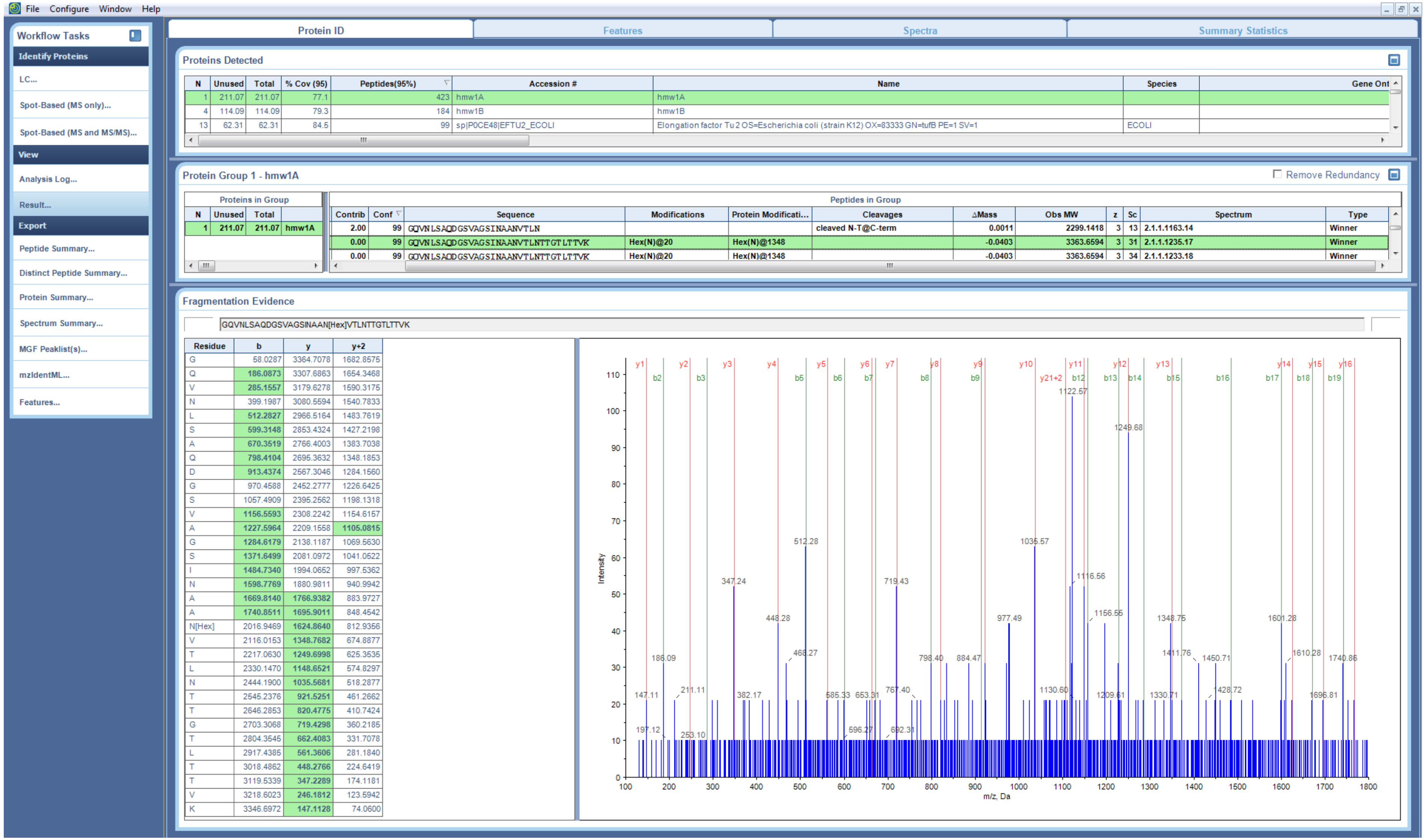


**Supplementary Figure S15. MS/MS peptide identification of GQVNLSAQDGSVAGSINAAN[+162.053]VTLNTTGTLTTVK**

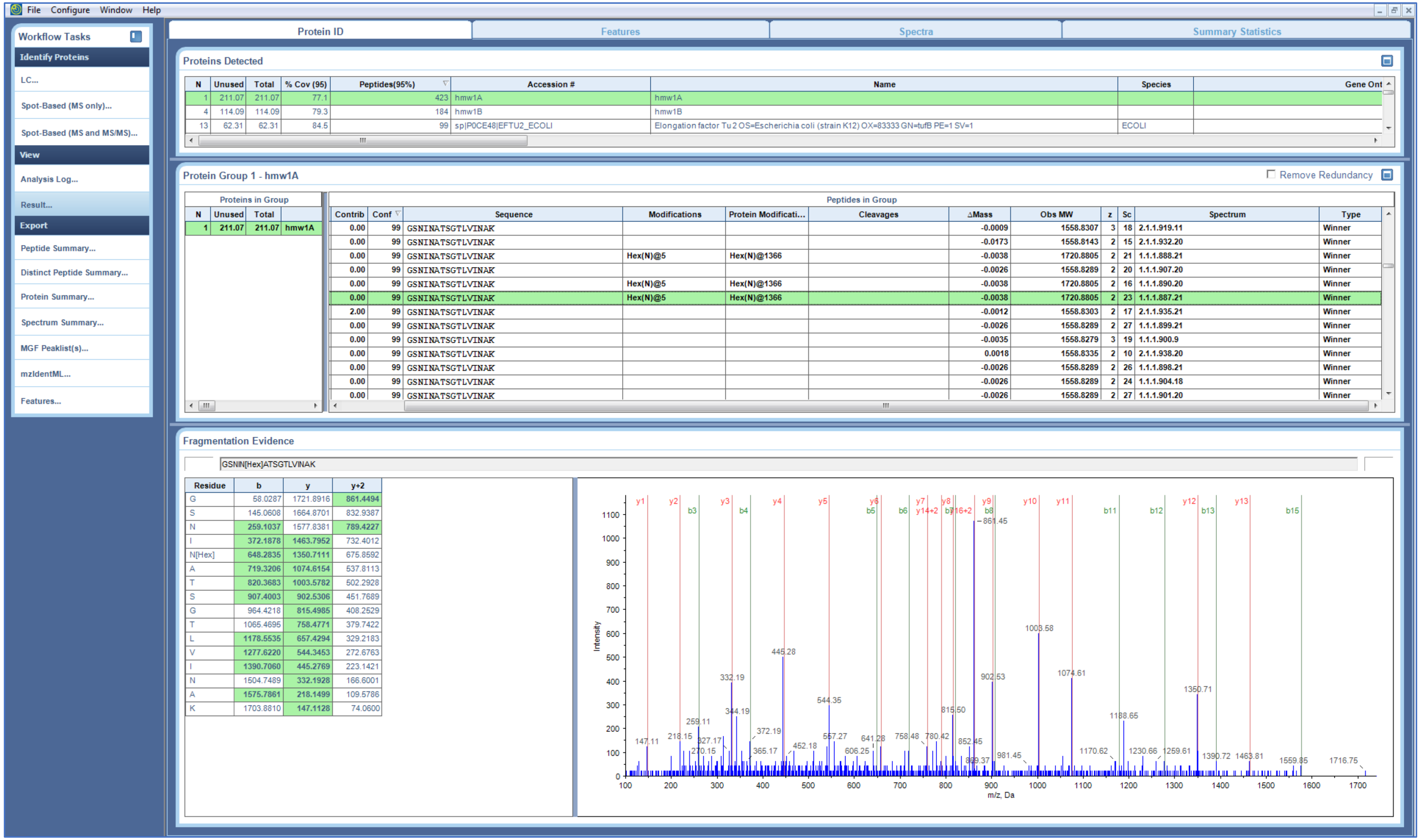

**Supplementary Figure S16. MS/MS peptide identification of GSNIN[+162.053]ATSGTLVINAK**

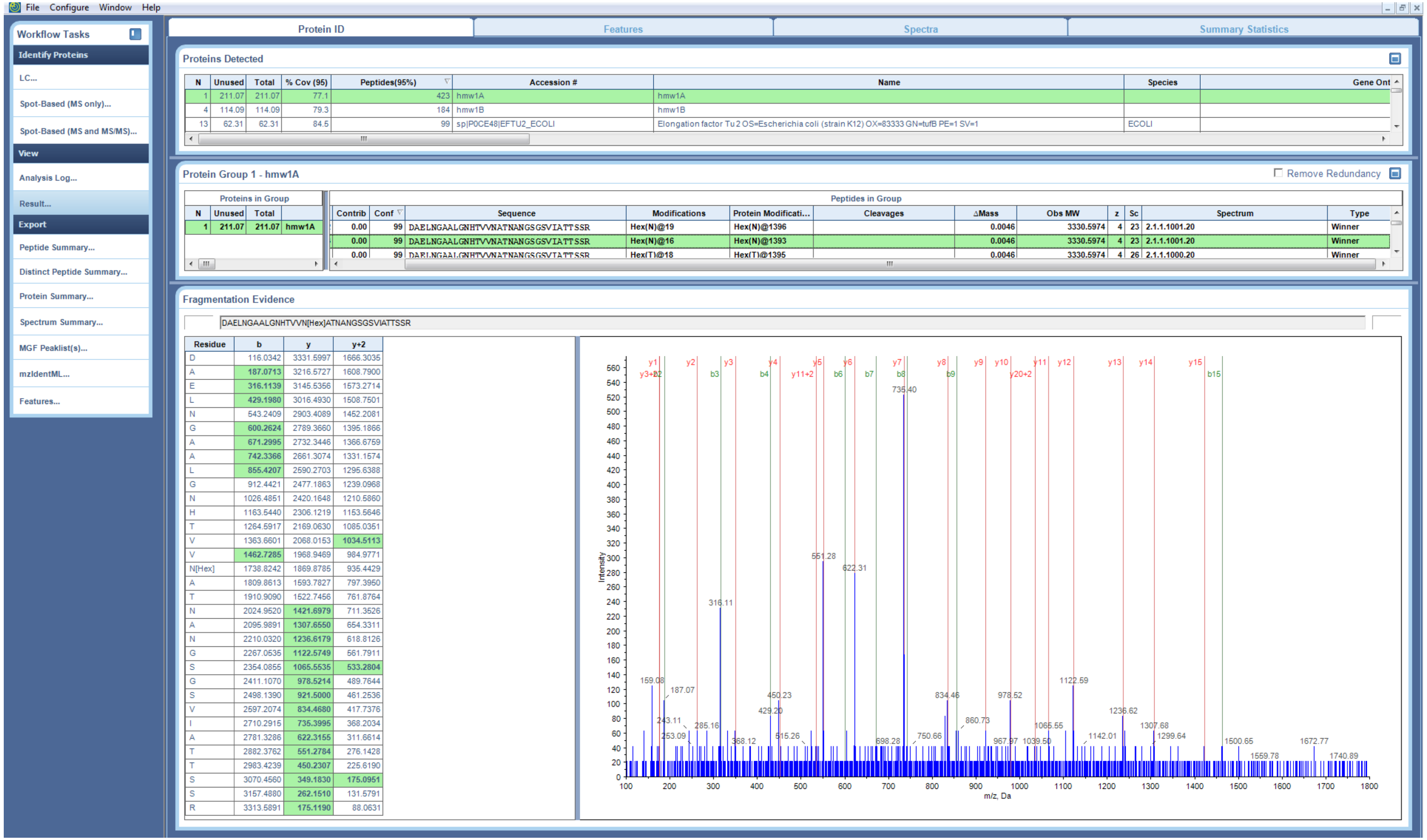


Supplementary Figure S17. MS/MS peptide identification of DAELNGAALGNHTVVN[+162.053]ATNANGSGSVIATTSSR

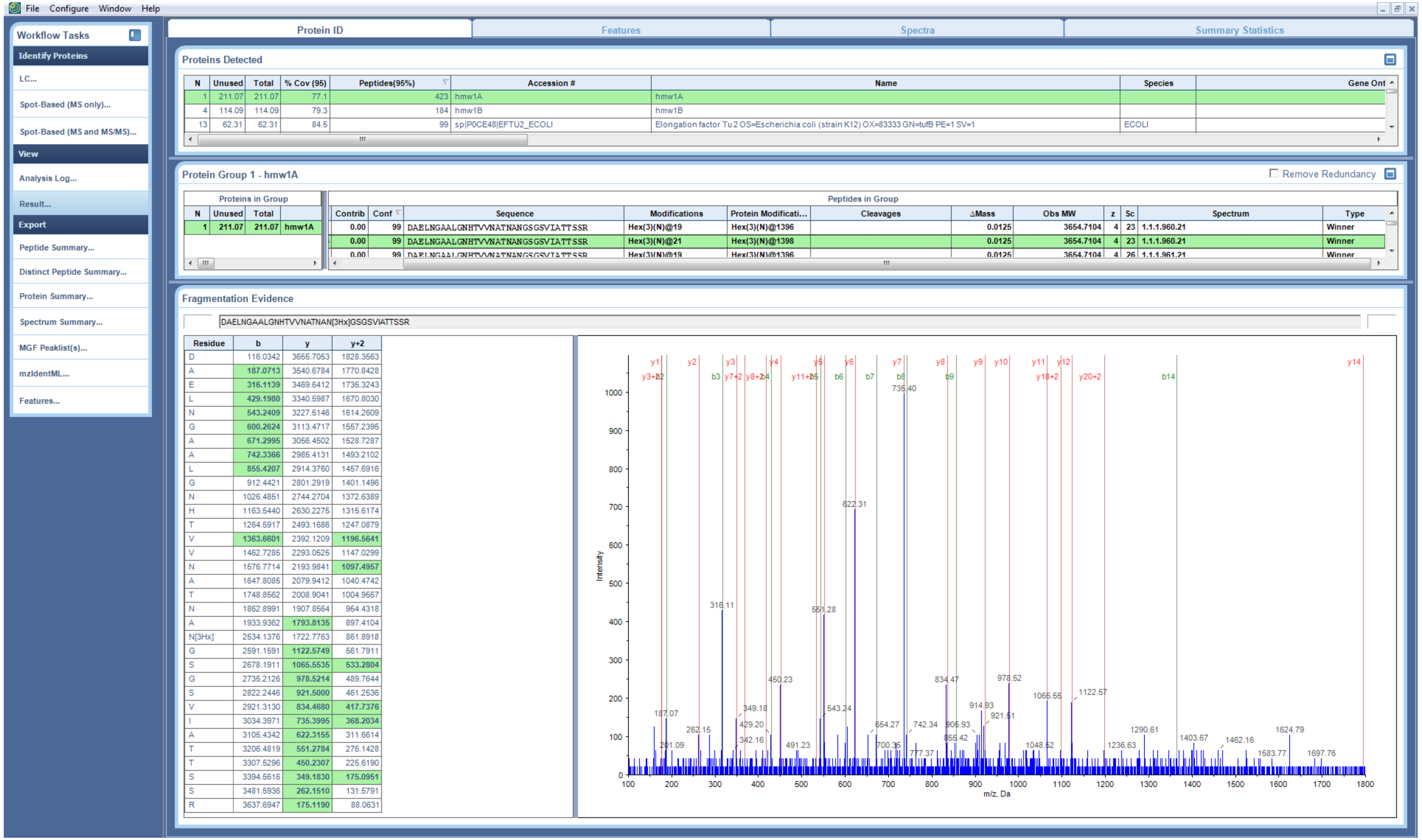


**Supplementary Figure S18. MS/MS peptide identification of DAELNGAALGNHTVVN[+486.159]ATNANGSGSVIATTSSR.** Site assignment of hexose-Asn is ambiguous, and assigned in Table 2 based on the presence of glycosylation sequons.

```
                   1                                                 50
Sequence           MNKIYRLKFSKRLNALVAVSELARGCDHSTEKGSEKPARMKVRHLALKPL
JPred Prediction   ---EEEEEE-----EEEEEEE---------------------HHHHHHHH
                   51                                                100
Sequence           SAMLLSLGVTSIPQSVLASGLQGMDVVHGTATMQVDGNKTIIRNSVDAII
JPred Prediction   HHHHHHHH---------------------------------------EEE
                   101                                               150
Sequence           NWKQFNIDQNEMVQFLQENNNSAVFNRVTSNQISQLKGILDSNGQVFLIN
JPred Prediction   E---------EEEEEE-----EEEEEEE------EEEEEEE---EEEEE-
                   151                                               200
Sequence           PNGITIGKDAIINTNGFTASTLDISNENIKARNFTFEQTKDKALAEIVNH
JPred Prediction   ---EEE-----E-----EEE--------------EE---------EEE--
                   201                                               250
Sequence           GLITVGKDGSVNLIGGKVKNEGVISVNGGSISLLAGQKITISDIINPTIT
JPred Prediction   ----------EEEE--------EEE-----EEEE----EEE---------
                   251                                               300
Sequence           YSIAAPENEAVNLGDIFAKGGNINVRAATIRNQGKLSADSVSKDKSGNIV
JPred Prediction   EEE-------------E-----EEE---------EEE-----------EE
                   301                                               350
Sequence           LSAKEGEAEIGGVISAQNQQAKGGKLMITGDKVTLKTGAVIDLSGKEGGE
JPred Prediction   EE---EEEEE--------------EEEEE---EEE----EEEE------E
                   351                                               400
Sequence           TYLGGDERGEGKNGIQLAKKTSLEKGSTINVSGKEKGGRAIVWGDIALID
JPred Prediction   EEEE----------------EEE---------------EEE-----EEE-
                   401                                               441
Sequence           GNINAQGSGDIAKTGGFVETSGHDLFIKDNAIVDAKEWLLD
JPred Prediction   --EEEE----------EEEEE---EEE----E---------
```

**Supplementary Figure 19. Secondary structure prediction of HMW1A signal peptide and pro-piece domains using JPred.**

```
                   442                                              491
Sequence           PDNVSINAETAGRSNTSEDDEYTGSGNSASTPKRNKEKTTLTNTTLESIL
JPred Prediction   ---EEE--------------------------------EEE---------
                   492                                              541
Sequence           KKGTFVNITANQRIYVNSSINLSNGSLTLWSEGRSGGGVEINNDITTGDD
JPred Prediction   -----EEEE----EEE---------EEEEE--------EE----------
                   542                                              591
Sequence           TRGANLTIYSGGWVDVHKNISLGAQGNINITAKQDIAFEKGSNQVITGQG
JPred Prediction   -------------EEE-----------EEE--------------EEE---
                   592                                              641
Sequence           TITSGNQKGFRFNNVSLNGTGSGLQFTTKRTNKYAITNKFEGTLNISGKV
JPred Prediction   EEEE-----EE---EEEE------EEE----------------EE-----
                   642                                              691
Sequence           NISMVLPKNESGYDKFKGRTYWNLTSLNVSESGEFNLTIDSRGSDSAGTL
JPred Prediction   EE----------------------EEE-----------------------
                   692                                              741
Sequence           TQPYNLNGISFNKDTTFNVERNARVNFDIKAPIGINKYSSLNYASFNGNI
JPred Prediction   ---------E------EEE-----EEE-----------------------
                   742                                              791
Sequence           SVSGGGSVDFTLLASSSNVQTPGVVINSKYFNVSTGSSLRFKTSGSTKTG
JPred Prediction   EE------EEE------------EEE----EE------EEEE--------
                   792                                              841
Sequence           FSIEKDLTLNATGGNITLLQVEGTDGMIGKGIVAKKNITFEGGNITFGSR
JPred Prediction   EE--------------E--------------------EEE----EEEE--
                   842                                              891
Sequence           KAVTEIEGNVTINNNANVTLIGSDFDNHQKPLTIKKDVIINSGNLTAGGN
JPred Prediction   ---------EEE-----EEE------------------------------
                   892                                              941
Sequence           IVNIAGNLTVESNANFKAITNFTFNVGGLFDNKGNSNISIAKGGARFKDI
JPred Prediction   --------------------------------------------------
                   942                                              991
Sequence           DNSKNLSITTNSSSTYRTIISGNITNKNGDLNITNEGSDTEMQIGGDVSQ
JPred Prediction   -------EE--------------------------------------EE-
                   992                                              1041
Sequence           KEGNLTISSDKINITKQITIKAGVDGENSDSDATNNANLTIKTKELKLTQ
JPred Prediction   ----EEE--------------------------------EE---------
                   1042                                             1091
Sequence           DLNISGFNKAEITAKDGSDLTIGNTNSADGTNAKKVTFNQVKDSKISADG
JPred Prediction   ----------EEE-------E-----------------------------
                   1092                                             1141
Sequence           HKVTLHSKVETSGSNNNTEDSSDNNAGLTIDAKNVTVNNNITSHKAVSIS
JPred Prediction   --EEE----------------------EEE--------------------
                   1142                                             1191
Sequence           ATSGEITTKTGTTINATTGNVEITAQTGSILGGIESSSGSVTLTATEGAL
JPred Prediction   --------------------------------------------------
                   1192                                             1241
Sequence           AVSNISGNTVTVTANSGALTTLAGSTIKGTESVTTSSQSGDIGGTISGGT
JPred Prediction   --------------------------------------------------
                   1242                                             1291
Sequence           VEVKATESLTTQSNSKIKATTGEANVTSATGTIGGTISGNTVNVTANAGD
JPred Prediction   EEE-----------------------------------------------
                   1292                                             1341
Sequence           LTVGNGAEINATEGAATLTTSSGKLTTEASSHITSAKGQVNLSAQDGSVA
JPred Prediction   --------EEE-----EE--------------------------------
                   1342                                             1391
Sequence           GSINAANVTLNTTGTLTTVKGSNINATSGTLVINAKDAELNGAALGNHTV
JPred Prediction   -------------------------------EE-----------------
                   1392                                             1441
Sequence           VNATNANGSGSVIATTSSRVNITGDLITINGLNIISKNGINTVLLKGVKI
JPred Prediction   --------------------------------EEE------EEEE----E
                   1442                                             1491
Sequence           DVKYIQPGIASVDEVIEAKRILEKVKDLSDEEREALAKLGVSAVRFIEPN
JPred Prediction   E--------------------------------------EEE--------
                   1492                                             1536
Sequence           NTITVDTQNEFATRPLSRIVISEGRACFSNSDGATVCVNIADNGR
JPred Prediction   ----------------EEEEE----EEE------EEEEEE-----
```

**Supplementary Figure 20. Secondary structure prediction of mature HMW1A using JPred.**
